# Potentiating the anti-tumor response of tumor infiltrated T cells by NAD^+^ supplementation

**DOI:** 10.1101/2020.03.21.001123

**Authors:** Yuetong Wang, Fei Wang, Lihua Wang, Shizhen Qiu, Yufeng Yao, Xuexue Xiong, Xuyong Chen, Quanquan Ji, Jian Cao, Dake Li, Liye Zhang, Ruoning Wang, Haopeng Wang, Gaofeng Fan

**Affiliations:** School of Life Science and Technology, ShanghaiTech University, Shanghai, China; Department of Gynecology, International Peace Maternity and Child Health Hospital, School of Medicine, Shanghai Jiao Tong University Shanghai, China; Center for Childhood Cancer and Blood Diseases, Hematology/Oncology & BMT, The Research Institute at Nationwide Children’s Hospital, Ohio State University, Columbus, OH, United States; State Key Laboratory of Cell Biology, Shanghai Institutes for Biological Sciences, Chinese Academy of Sciences, Shanghai 200031, China; Department of Gynecology, Women’s Hospital of Nanjing Medical University, Nanjing Maternity and Child Health Care Hospital, Nanjing, China

**Author notes:** Equal Contribution (Yuetong Wang, Fei Wang, Lihua Wang, Shizhen Qiu).

## Abstract

Tumor immunotherapies have provided clinical benefits, yet great potential remains for optimizing therapeutic effects. Here, we show that low NAD^+^ levels restrict the function of tumor infiltrating T lymphocytes (TILs). TILs harvested from human ovarian tumor tissues showed decreased NAD^+^ levels compared with T cells from paired peripheral blood samples. The combination of whole-genome CRISPR and large-scale metabolic inhibitor screens implicated the NAD^+^ biosynthesis enzyme nicotinamide phosphoribosyltransferase (NAMPT) is required for T cell activation. Further isotopic labeling and LC-MS studies confirmed that NAD^+^ depletion suppressed mitochondrial energy biosynthesis in T cells. Excitingly, NAD^+^ supplementation significantly enhanced the tumor cell-killing efficacy of CAR-T cells *ex vivo*, and extended animal survive in both adoptive CAR-T model and immune checkpoint blockade treatment models *in vivo*. This study demonstrates an over-the-counter nutrient supplement NAD^+^ could robustly boost the efficacy of T cell-based immunotherapy and provides insights into the cellular basis of T cell metabolic reprogramming in treating cancers.

**One Sentence Summary:** NAD^+^ supplementation during cancer immunotherapies significantly enhances T cell activation and tumor killing capacity.

## Main Text

Cancer immunotherapies including adoptive transfer of naturally-occurring tumor infiltrating lymphocytes (TIL) and genetically-engineered T cells, as well as the use of immune checkpoint inhibitors to boost the function of T cells have emerged as promising approaches to achieve durable clinical responses of otherwise treatment-refractory cancers ^1-5^. Although cancer immunotherapies have been successfully utilized in the clinic for subsets of patients, there are several limitations which prevent the broad use of these therapies for entire patient populations ^6,7^. Given the function of T cells as key mediators for tumor destruction, their characteristics (e.g., durability, longevity, and killing efficiency, etc.) substantially determine the clinical outcomes of many immunotherapies ^8-11^. Studies have established that successful clearance of tumors mediated by infiltrated T cells can be limited by physical barriers generated by stroma cells ^12^, immuno-suppressive networks ^13^, and nutrient limitations within the microenvironment ^14,15^. Thus, efforts to promote the stemness, proliferation and activation capacity of T cells should enable improvements to the efficacy of cancer immunotherapies.

Recently, cellular metabolic processes have been reported to shape T cell differentiation and functional activity ^16-18^. Modulation of metabolic process such as fatty acid catabolism can improve T cell activation and therapeutic function ^19-24^. It was also reported that there is a strong link between metabolic activity in tumor infiltrated T cells (TIL) and their effector function ^25,26^, and stimulation mitochondrial biogenesis via enforced expression of PGC1α resulted in superior anti-tumor function of T cells ^23^. These studies reinforce that efforts on investigating immune response and metabolic regulations required for TILs when combating with tumors will help us to understand the bio-energetic requirements of T cell function and help in the optimization of tumor immunotherapies.

Seeking to identify genes involved in the regulation of T cell activation, we performed a genome-wide CRISPR screen which containing over 250,000 total sgRNA using the Jurkat T cell line by adopting a previously reported sgRNA-based strategy ^27^. Subsequently, we analyzed our Jurkat T cell screening data alongside the previously reported primary human T cell data ^28^ and identified the top 50 target gene hits common to both screens (Fig. 1a green dots, Extended Data Fig. 1a, and Supplementary Table 1). In concert with this genetic screen, and aiming to identify the metabolic pathways that are essential for T cell activation, we also undertook chemical screens of both Jurkat T cell and primary human T cells using the 199 compounds of the Metabolism Compound Library (Selleck, L5700) and monitoring T cell activation with multiple markers (Fig. 1b-d and Extended Data Fig. 1b).

**Fig.1.**
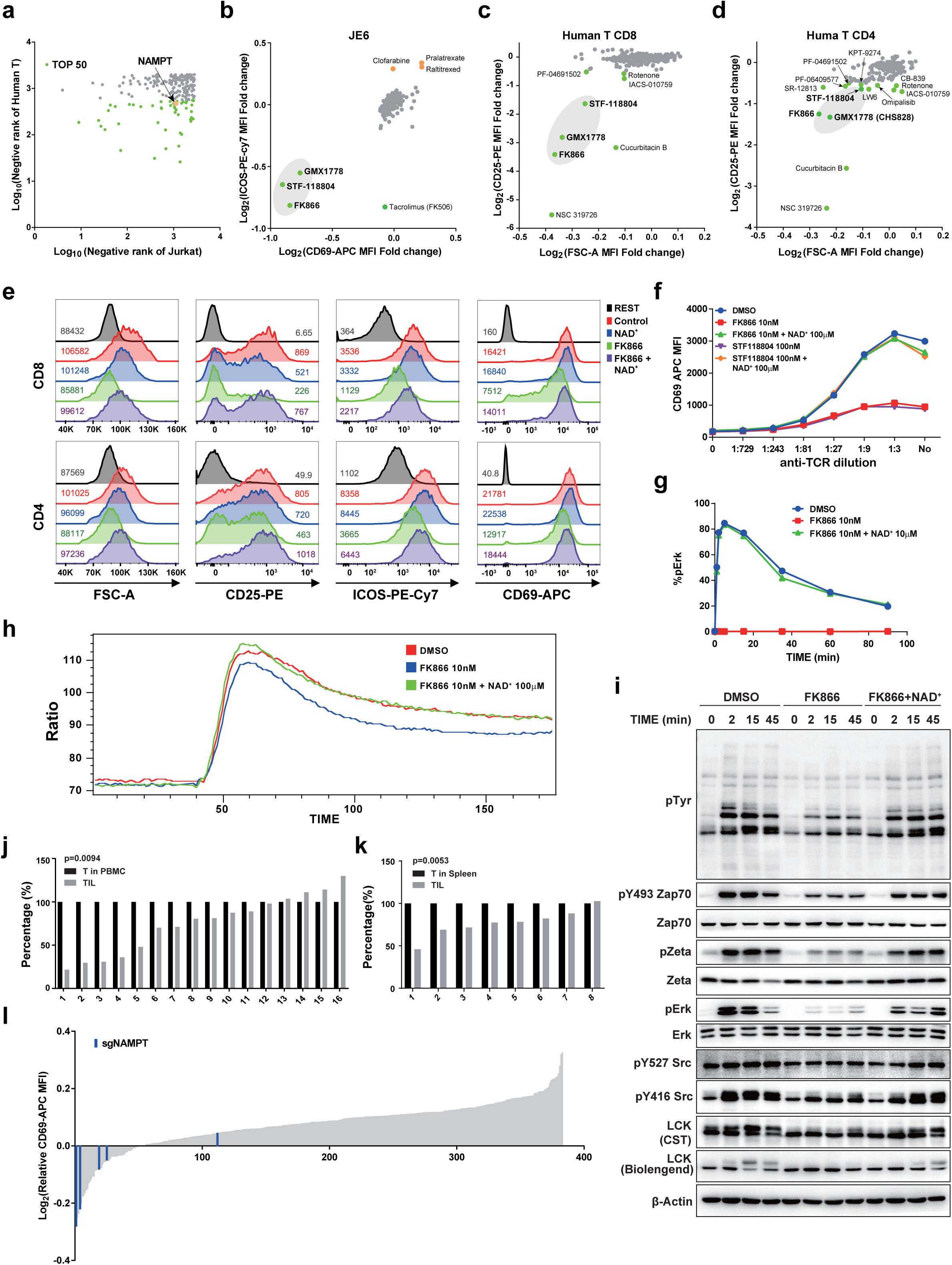
NAMPT-mediated NAD^+^ production is essential for T cell activation. **(a)** Whole genome sgRNA screens were performed in both Jurkat and primary human T cells ^28^ to identify genes involved in T cell activation. The top 50 target gene screening hits common to both cell types (green dots). The orange dot represents the *NAMPT* gene. **(b)** Metabolic inhibitor screening in Jurkat (JE6) cells. Jurkat cells were pre-treated with a panel of 199 inhibitors (10nM) for 24 hours, followed by anti-TCR stimulation for 16 hours. Cells were then stained with anti-CD69-APC and anti-ICOS-PE-Cy7 for 40min on ice. The surface levels of these markers were measured by flow cytometry. **(c-d)** Metabolic inhibitor screening in primary human T cells. Isolated human CD8^+^ T cells (c) and CD4^+^ T cells (d) were treated with the same panel of inhibitors (100nM) for 24 hours, followed by transfer into well of 96-well plates pre-coated with anti-CD3 and anti-CD28 antibodies and cultured for 16 hours. After stimulation, cells were stained with anti-CD69-APC, anti-CD25-PE, and anti-ICOS-PE-Cy7 for 40min on ice. The surface levels of these markers were measured based on immunofluorescence. **(e)** Isolated human T cells among PBMCs (both CD8^+^ and CD4^+^) were pre-treated as indicated (NAD^**+**^, FK866 or FK866 + NAD^**+**^) for 24 hours. Cells were then transferred to a plate coated with anti-CD3 and anti-CD28 for stimulation. Cell volumes and cell-surface expression levels of CD25, ICOS, and CD69 were measured using flow cytometry, respectively. **(f)** Cell surface expression level of CD69 in anti-TCR activated Jurkat cells cultured with the NAD^+^ synthesis inhibitors (FK866 or STF118804) or with NAD^+^; Jurkat cells were first treated with inhibitors or NAD^+^ for 24 hours, followed by stimulation with anti-TCR for 16 hours. **(g)** Monitoring of the ERK phosphorylation level in Jurkat cells; Jurkat cells were treated as indicated (DMSO, FK866, or FK866 + NAD^+^) for 24 hours followed by resting in RPMI1640 medium without FBS for 1 hour. Then cells were further stimulated with anti-TCR antibody and collected at the indicated time points and stained with a phospho-ERK specific antibody. **(h)** Jurkat cells were treated as indicated (DMSO, FK866 or FK866 + NAD^**+**^) for 24 hours, followed by Calcium probe Indo-1 labeling. Cells were pre-incubated with 2× anti-TCR antibody and 2 μM Ionomycin at 37°C for 5 minutes prior to measurement by flow cytometry|. **(i)** Immunoblot analysis of the levels of phosphotyrosine, phospho-ZAP70, phospho-ERK, phospho-SRC, phospho-LCK, phospho-Zeta in Jurkat cells stimulated with anti-TCR antibody. Jurkat cells were treated as indicated (DMSO, FK866 or FK866 + NAD^**+**^) for 24 hours, followed by resting in RPMI1640 medium without FBS for 1 hour. Then cells were further stimulated with anti-TCR antibody and collected at the indicated time points for immunoblotting. **(j to k)** NAD^+^ concentration measurement in paired TILs and T cells among PBMCs of ovarian cancer patient samples (j), and paired TILs and T cells in spleen of melanoma mouse allograft tumors (k). For TILs and T in PBMCs of ovarian cancer patient samples, samples were stained with anti-human CD3 antibody and isolated by FACS. For mouse samples, 6-week-old B6 mice were subcutaneously transplanted with 1×10^6^ B16F10 mouse melanoma cancer cells. Two weeks later, both tumors and spleens were harvested and analyzed by sorting for, respectively, CD3^+^ tumor infiltrated (TIL) cells and spleen CD3^+^ cells. **(l)** sgRNA screening of NAD^+^ metabolism related genes apparently required for T cell activation. JX003 cells (a single clone selected from Jurkat cells stably expressing Cas9) were first infected with a lentivirus expressing one of 390 sgRNAs targeting each previously annotated NAD^+^ metabolism related gene (94 genes, 4∼5 sgRNA/gene, Table S3). The viral titer used for infection aimed to infect half of the Jurkat cell population (MOI=0.5). After seven days, cells were stimulated with anti-TCR antibody, and changes in the expression level of CD69 were calculated as follows: the CD69 luminescence intensity of infected Jurkat cells divided by the CD69 luminescence intensity of uninfected Jurkat cells.

Combined analysis of the genetic and metabolic screen data revealed the conspicuous commonality of the NAD^+^ biosynthesis enzyme nicotinamide phosphoribosyltransferase (NAMPT) (Fig. 1a-d and Extended Data Fig. 1a-b). Specifically, the *NAMPT* gene identified by both of the sgRNA CRISPR screens, and three of the small molecule compounds from the library known to target NAMPT (FK866, STF-118804, and GMX1178) ranked among the most disruptive compounds to T cell activation. These analyses also revealed that all three of the NAMPT inhibitors apparently had more pronounced effects on activation of CD8^+^ T cells than on CD4^+^ T cell activation (Fig. 1c-d and Extended Data Fig.S1b).

To confirm the essential role of NAD^+^ in T cell activation, we treated anti-CD3 and anti-CD28-stimulated human primary T cells with the NAD^+^ synthesis inhibitor FK866. The stimulatory effects were remarkably repressed by inhibition of NAD^+^ biosynthesis: FK866 treatment drastically reduced the up-regulation of T cell activation markers such as CD69, CD25, and ICOS, and apparently blocked the increase of cell size of activated T cells (Fig.1e and Extended Data Fig. 1c). We observed similar effects upon FK866 treatment of expanded primary T cells (Extended Data Fig. 1d-e, and 1g). And, we found that the intracellular NAD^+^ level could be significantly increased by simply adding NAD^+^ into culture medium (Extended Data Fig. 1d), suggesting direct cellular NAD^+^ uptake in T cells, which has also been reported in macrophages, neuronal cell and hepatoma cells ^29,30^. Further, assays with Jurkat cells showed that the NAMPT inhibitors FK866 and STF-118804 suppressed cell activation upon anti-TCR stimulation in a dose-dependent manner (Extended Data Fig. 1f, 1j-m), and this suppression was achieved in multiple stimulation contexts (Fig. 1f and Extended Data Fig. 1h).

We next examined the effects of these inhibitors on T cell signaling events, including calcium flux and phosphorylation of T cell signaling proteins such as ZAP70, LCK, and ERK. FK866 treatment significantly decreased both calcium flux and the phosphorylation levels of T cell signaling proteins in stimulated Jurkat cells (Fig.1g-i, and Extended Data Fig. 1i). Importantly, this potent repression could be rescued by adding exogenous NAD^+^ in the cell culture medium (Fig.1e-i, Extended Data Fig. 1c-i, 1k, and 1m). These results together establish that NAD^+^ is required for T cell activation.

Previous studies have reported that metabolic stress can impair the function of TILs ^24^ and can also limit their reinvigoration capacity ^31^. We therefore examined intracellular NAD^+^ levels in TILs by analyzing paired T cells from peripheral blood mononuclear cells (PBMCs) and TILs from 16 ovarian cancer patients. We also analyzed paired T cells from spleen and TILs from 8 melanoma allograft mice. In both sample types, the NAD^+^ levels were significantly lower in the respective TILs as compared to paired T cells in patients’ PBMCs and to paired T cells in murine spleen, separately (Fig.1j-k). Since it was reported that NAD^+^ level was higher in activated T cells ^32^, we concluded that the decreased NAD^+^ level in tumor infiltrated T cells was caused by tumor microenvironment instead of T cell activation state. We also analyzed the differentially expressed genes between TILs and T cells in PBMCs from a mouse model of autochthonous melanoma, which revealed enrichment for the nicotinate and nicotinamide metabolism KEGG pathway (Extended Data Fig. 2a). These results suggest that decreased NAD^+^ levels in TILs may contribute to the previously reported decreases in the killing capacity and reinvigoration capacity of TILs under metabolic stress.

NAD^+^ fluxes have been reported to vary widely across different cell types and tissues, and accordingly, different NAD^+^ synthesis pathways, which feature specialized enzymes, are adopted by each cell type ^33,34^. To evaluate the major NAD^+^ synthesis pathway in T cells, we conducted an sgRNA screen in anti-TCR stimulated Jurkat T cells for 94 genes with NAD^+^-related functions concatenated in a previous review paper ^35^. This again revealed that NAMPT is essential for T cell activation (Fig. 1l, and Extended Data Fig. 2b-c, and 2e), and subsequent experiments confirmed that knockdown of *NAMPT* decreased the NAD^+^ level by more than 50%, illustrated the salvage NAD^+^ synthesis pathway as the main NAD^+^ source in T cells (Extended Data Fig. 2d).

Enzymes of the Sirtuin and PARP families which used NAD^+^ as a substrate are known to exert functions in, respectively, epigenetic regulation and in DNA damage repair ^36,37^. However, we found that inhibition of these enzymes with either sgRNA knockout or pharmacological inhibitors caused no significant changes to Jurkat T cells activation (Fig.1l and Extended Data Fig. 3a-d), suggesting that the NAD^+^’s role as a substrate for using in epigenetic regulation and DNA damage repair processes may be less relevant to T cell activation than its role as a cofactor in metabolic process.

To further assess NAD^+^ mediated regulation of T cell activation, we performed LC-MS based metabolite profiling of FK866-treated and control Jurkat T cells. We found that such inhibition of NAD^+^ synthesis resulted in significantly altered levels of metabolites of the glutaminolysis, glycolysis, and citric acid cycle pathways, findings indicative of decreased mitochondrial oxidative phosphorylation (Fig. 2a-b). We also repeated the LC-MS experiment with human PBMC cells, and glycolysis and citric acid cycle pathway were also decreased in FK866 treated PBMC cells (Extended Data Fig. 4a-c), indicating NAD^+^ metabolism regulates mitochondrial oxidative phosphorylation in both Jurkat T cells and PBMC cells. Further isotopic labeling experiments using ^3^H-3-Glucose and ^14^C-U-Glutamine confirmed the inhibitory effect of FK866 treatment on glycolysis and glutaminolysis (Fig. 2c-d). Consistent with a NAMPT-mediated regulation of mitochondrial oxidative phosphorylation, FK866 treatment of Jurkat T cells caused significant decreases in lactate, citric acid, succinic acid, oxoglutaric acid, as well as increases in glucose and glutamine (Extended Data Fig. 5). Since the majority of ATP is generated in the TCA cycle ^38^, we propose that NAD^+^ levels apparently regulate the energy generation functions of mitochondria in T cells. Transmission electron microscopy (TEM) showed that depriving Jurkat T cells or primary T cells of NAD^+^ (either by inhibitor treatment or by knocking down *NAMPT*) significantly decreased the number of mitochondria and reduced the maximum width of cristae (Fig. 2e-f, and Extended Data Fig. 6a-c, 6f). And mitochondrial oxygen consumption rate measurements in Jurkat T cells or primary T cells cells showed that inhibition of NAD^+^ biosynthesis weakened the respiratory capacity of mitochondria (Fig. 2g, and Extended Data Fig. 6d, 6g). Consequently, the total cellular level of ATP was also decreased in NAD^+^ deprived cells (Fig. 2h-i, and Extended Data Fig. 6e).

**Fig.2.**
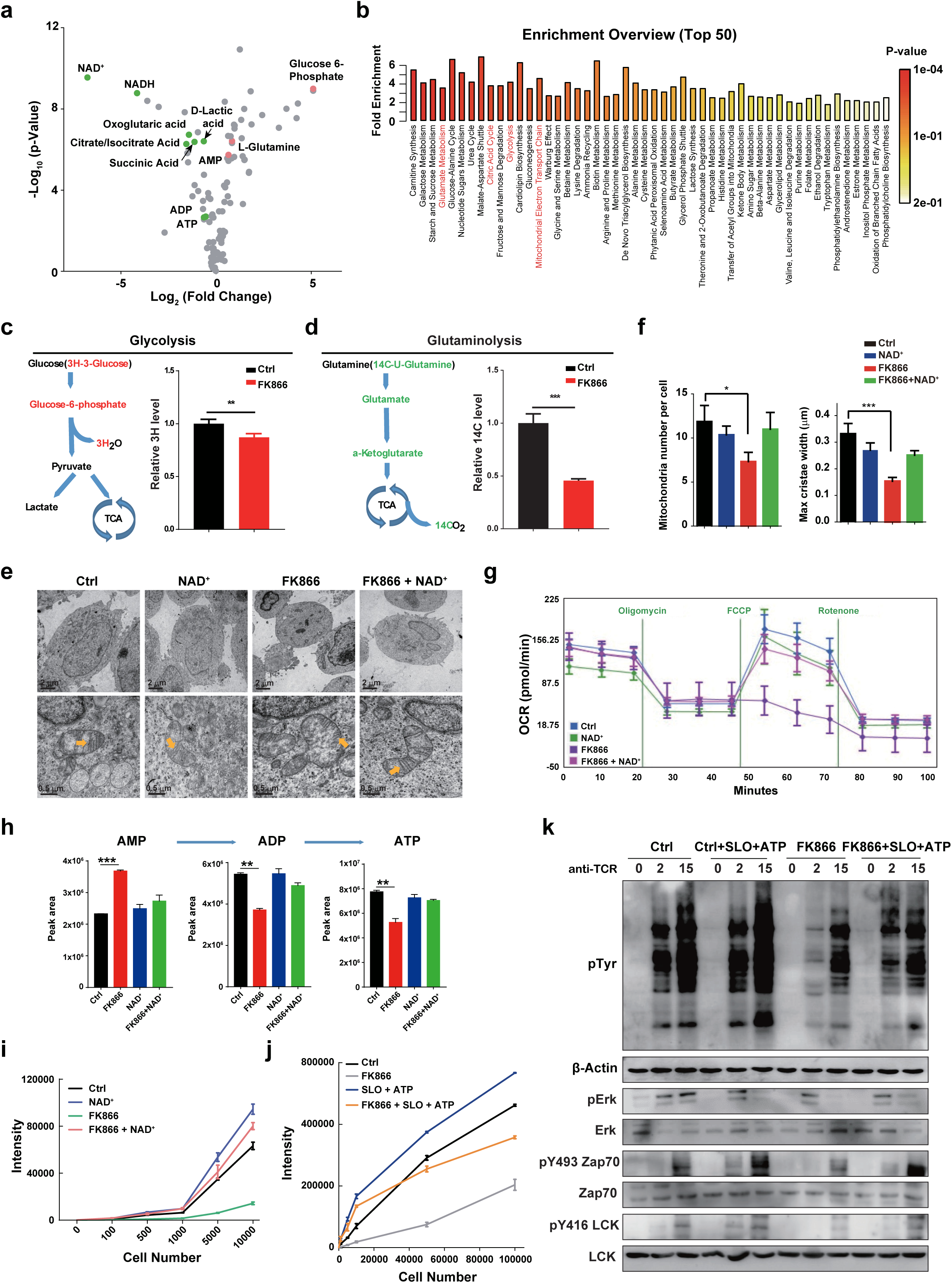
NAD^+^ regulates cellular energy metabolism through the TCA cycle in T cells. **(a)** LC-MS based metabolic profiling of Jurkat cells treated with vehicle or FK866. 1*10^7^ Jurkat cells, with or without 10nM FK866 co-treatment for 24 hours. Each group comprised six replicates. The green dots represent the metabolites down-regulated by FK866 treatment, and the red dots represent metabolites up-regulated by FK866 treatment. **(b)** Enrichment analysis of significantly changed metabolites. All significantly changed metabolites were used for metabolic pathway analysis with the MetaboAnalyst 4.0. **(c to d)** Isotopic labeling with ^3^H-3-glucose (c) and ^14^C-U-glutamin (d) in Jurkat cells. ** p < 0.01, *** p < 0.001. Jurkat cells were treated with FK866 for 24 hours prior to the isotopic labeling experiments. **(e to f)** Representative transmission electron microscope (TEM) images showing the number of, and morphological changes in, mitochondria in Jurkat cells upon treatment (e) with inferential statistical analysis using t tests (f). * p < 0.05, *** p < 0.001. Jurkat cells were treated as indicated (Ctrl, NAD^**+**^, FK866 or FK866 + NAD^**+**^) for 24 hours. Then cells were harvested and prepared for TEM. **(g)** Oxygen consumption rate (OCR) under the basal condition and in response to the indicated mitochondrial inhibitors. Jurkat cells were treated as indicated (Ctrl, NAD^**+**^, FK866 or FK866 + NAD^**+**^) for 24 hours. **(h)** ATP/ADP/AMP levels detected by LC-MS in Jurkat cells. ** p < 0.01, *** p < 0.001. 1×10^7^ Jurkat cells were treated as indicated for 24 hours. Each sample group comprised six replicates. **(i)** Cellular ATP level measured by CellTiter-Glo assay in Jurkat cells treated as indicated (Ctrl, NAD^**+**^, FK866 or FK866 + NAD^**+**^) for 24 hours. **(j)** Cellular ATP level detected by CellTiter-Glo assay in Jurkat cells treated with FK866 or streptolysin O (SLO) and ATP. Cells were treated with or without FK866 for 24 hours. Then, cells were treated with SLO for 1 hour, followed by 10min of incubation with an ATP-containing solution. Cells were thoroughly washed with PBS before the ATP concentration measurement. **(k)** Immunoblot analysis of the levels of phosphotyrosine, phospho-ZAP70, phospho-ERK, phospho-LCK in Jurkat cells stimulated with anti-TCR and with or without FK866 treatment for 24 hours. Cells were then treated with SLO for 1 hour, followed by stimulation with anti-TCR and treatment with ATP for the indicated times.

Although these results link T cell activation and intracellular ATP levels, we wanted to determine if there is a direct causal relationship. To this end, we used various strategies to manipulate glucose catabolism, glutamine catabolism, and intracellular ATP levels (Extended Data Fig. 7). Culturing Jurkat T cells with glucose or glutamine free medium (Extended Data Fig. 7a, 7c), and incubation of Jurkat T cells with glucose catabolism inhibitor (2-DG) or glutamine catabolism inhibitor (CB-839) (Extended Data Fig. 7b, 7d) could reduce Jurkat T cell activation. Moreover, restoration of cellular ATP levels rescued the activation capacity in all of these variously cultured cells (Fig. 2j-k, and Extended Data Fig. 7e-l). Collectively, these results support that NAD^+^ modulates T cell activation by regulating mitochondrial energy production.

Our findings prompted us to further examine the effect(s) of NAD^+^ supplementation on CAR-T cell function in solid tumors. Notably, recent studies have reported that NAD^+^ precursor supplementation is an effective strategy to protect against tissue aging and to lengthen life spans ^39-41^. We first confirmed the role of intracellular NAD^+^ in the anti-tumor activity of CAR-T cells. FK866 treatment significantly reduced the killing ability of anti-CD19-41BB CAR-T cells *ex vivo* without causing significant cell death in CAR-T cells (Fig. 3a, and Extended Data Fig. 8). Consistently, FK866 treatment significantly reduced cytokine production by stimulated CAR-T cells, including GzmB (Fig. 3b), IFN-γ (Fig. 3c), and IL-2 (Fig. 3d). To further study the NAD^+^ supplementation on T cell function, we overexpressed wild type or enzymatic dead mutant form of NAMPT (D313A) ^42^ in anti-CD19-41BB CAR-T cells and Jurkat T cells (Extended Data Fig. 9a, 9d). In both cells, overexpression of wild type NAMPT, but not NAMPT-D313A, could significantly increase the level of intracellular NAD^+^ (Extended Data Fig. 9b, 9e). And we found that overexpression of NAMPT could significantly stimulate the activity of Jurkat cells, especially when treated with FK866 (Extended Data Fig. 9c). In CAR-T cells, increase NAD^+^ level via overexpression of NAMPT could enhance tumor cell killing ability and elevate cytokine production (Extended Data Fig. 9f-g).

**Fig.3.**
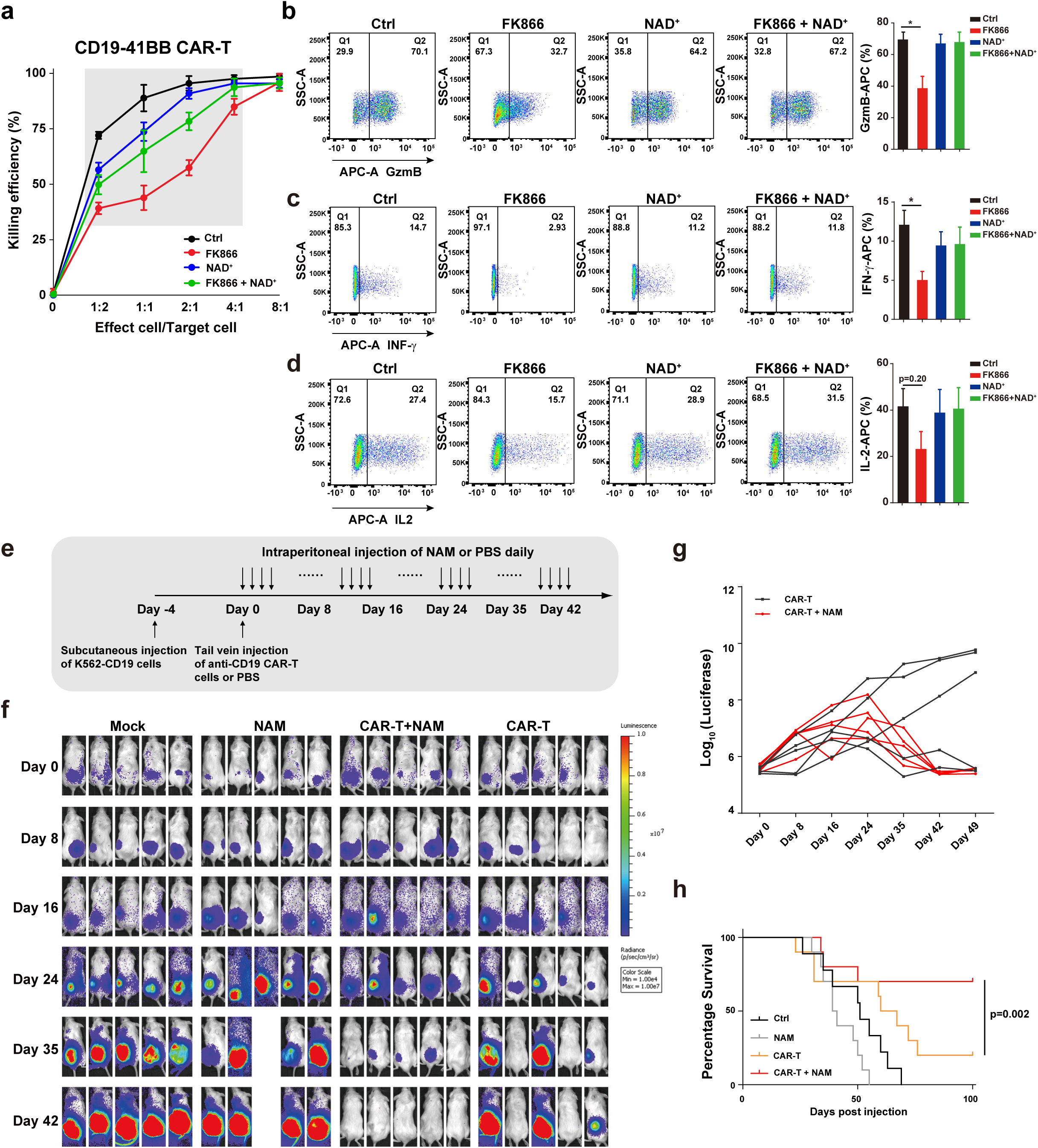
NAD^+^ supplementation enhances tumor killing function of CAR-T cells. **(a)** *In vitro* killing assay of target tumor K562-CD19 cells expressing a mCherry reporter; this is a chronic myelogenous leukemia cell line that overexpresses CD19. Anti-CD19-41BB CAR-T cells were pre-treated as indicated (Ctrl, NAD^**+**^, FK866 or FK866 + NAD^**+**^) for 48 hours, followed by co-culturing with mixed K562-CD19-mCherry and K562-WT (at a 1:1 ratio) cells at different ratios for 24 hours. The killing efficiency was monitored by flow cytometry and calculated as the cell death ratio of the K562-CD19-mCherry cells. **(b to d)** Cytokine production, including GzmB (b), IFN-γ (c), and IL-2 (d) of CAR-T cells. ** p < 0.01. CAR-T cells were pre-treated as indicated (Ctrl, NAD^**+**^, FK866 or FK866 + NAD^**+**^) for 48 hours, followed by co-culturing with K562-CD19 cells at a 1:1 ratio for 20 hours. Prior to fixation and staining, cells were treated with the intracellular protein transport inhibitor Brefeldin A for 4 hours. **(e)** Diagram of the experimental strategy used in the *in vivo* CAR-T cell killing assays. **(f to g)** Ability of adoptively transferred anti-CD19-41BB CAR-T cells to control the growth of s.c.-established K562-CD19-Luciferace tumors in NSG mice. Tumor growth was measured by bioluminescence imaging (f) and analyzed quantitatively (g). n=5. **(h)** Log-rank test of survival curves. Ctrl group n=9, NAM group n=10, CAR-T group n=10, and CAR-T+NAM group n=10.

To study the effects of NAD^+^ supplementation on immunotherapy *in vivo*, we first adopted a mouse model of CAR-T therapy (Fig. 3e). Briefly, mice were firstly subcutaneously inoculated with K562-CD19 tumor cells. Anti-CD19-41BB CAR-T cells were then adoptively transferred 4 days later via tail vein injection. To better explore anti-tumor function of NAD^+^ supplementation *in vivo*, CAR T cell doses were deliberately lowered to 10^6^ cells per mouse so that CAR-T had suboptimal anti-tumor potency. For the NAD^+^ supplementation, we compared several NAD^+^ precursor compounds before eventually choosing nicotinamide (NAM); NAM treatment caused comparable increases in intracellular NAD^+^ concentrations with direct NAD^+^ treatment which implied the comparable therapeutic effect of NAM and NAD^+^ (Extended Data Fig. 10a). Delivery of NAM by intraperitoneal injection after CAR-T cell transfer significantly increased the intracellular level of NAD^+^ in CAR-T cells (Extended Data Fig. 10b). Notably, NAM supplementation alone did not affect tumor growth (Extended Data Fig. 10c). We then explored the effects of NAM supplementation on T cells by comparing tumor killing efficiency of adopted CAR-T cells between NAM-treated and untreated mice. Remarkably, mice of the CAR-T plus NAM supplementation group showed better responses to the immunotherapy: these animals were all tumor-free by the end of the experiment (Fig. 3f-g, and Extended Data Fig. 10d-e). NAM supplementation also significantly extended the survive time of mice (Fig. 3h). Immunostaining of xenograft tumor tissues sections also showed that there were more TILs in the NAM-treated tumors (Extended Data Fig. 10f). Taken together, these results establish that supplementation with the NAD^+^ precursor NAM enhances CAR-T cell tumor cell killing function and improves the therapeutic efficacy of immunotherapy.

Immune-checkpoint blockade (ICB) strategy-derived antibodies that block cytotoxic T lymphocyte associated protein 4 (CTLA4), and programmed death 1 (PD-1) or its ligand PD-L1 have demonstrated unprecedented therapeutic efficacy in clinical trials, including metastatic melanoma and non-small cell lung cancer ^43-45^. However, there are still a large fraction of patients who have shown de novo and/or acquired resistance to ICB, probably due to cancer immunoediting ^46^, infiltrated T cell dysfunction 47, and tumor microenvironment effect 48. To bolster our observation further, we adopted immune-checkpoint blockade (ICB) animal model in combination with NAD^+^ supplementation. Briefly, B6 mice were firstly subcutaneously inoculated with B16F10 cells which were reported refractory to anti-PD-1 treatment to better evaluate the function of NAM supplementation. Then, mice were treated with NAM, anti-PD-1 or combined. Anti-PD-1 treatment or NAM supplementation per se could mildly inhibited tumor growth (Fig. 4a-d). Notably, mice of the anti-PD-1 plus NAM supplementation group showed significantly better responses to the immunotherapy: the tumor growth was effectively inhibited and 1 of the animals was almost cured (Fig. 4a-d). NAM supplementation also significantly extended the survive time of mice (Fig. 4e). We also examined the percentage of tumor infiltrated T cells (TILs) by anti-CD3 staining of tumors, and found that anti-PD-1 treatment or NAM supplementation per se could significantly increase the percentage of TILs (Fig. 4f-g). Mice with combination treatment showed the highest percentage of TILs, which indicated the strongest immune response (Fig. 4f-g). We also used MC38 tumor cell model to study the function of NAD^+^ supplementation in conjunction with anti-CTLA-4 treatment. Again, mice with combination treatment of anti-CTLA-4 treatment and NAM supplementation showed the slowest tumor growth and longest survive (Extended Data Fig. 11a-c). In summary, NAD^+^ supplementation could significantly enhance immunotherapeutic effect of ICB in solid tumor models.

**Fig.4.**
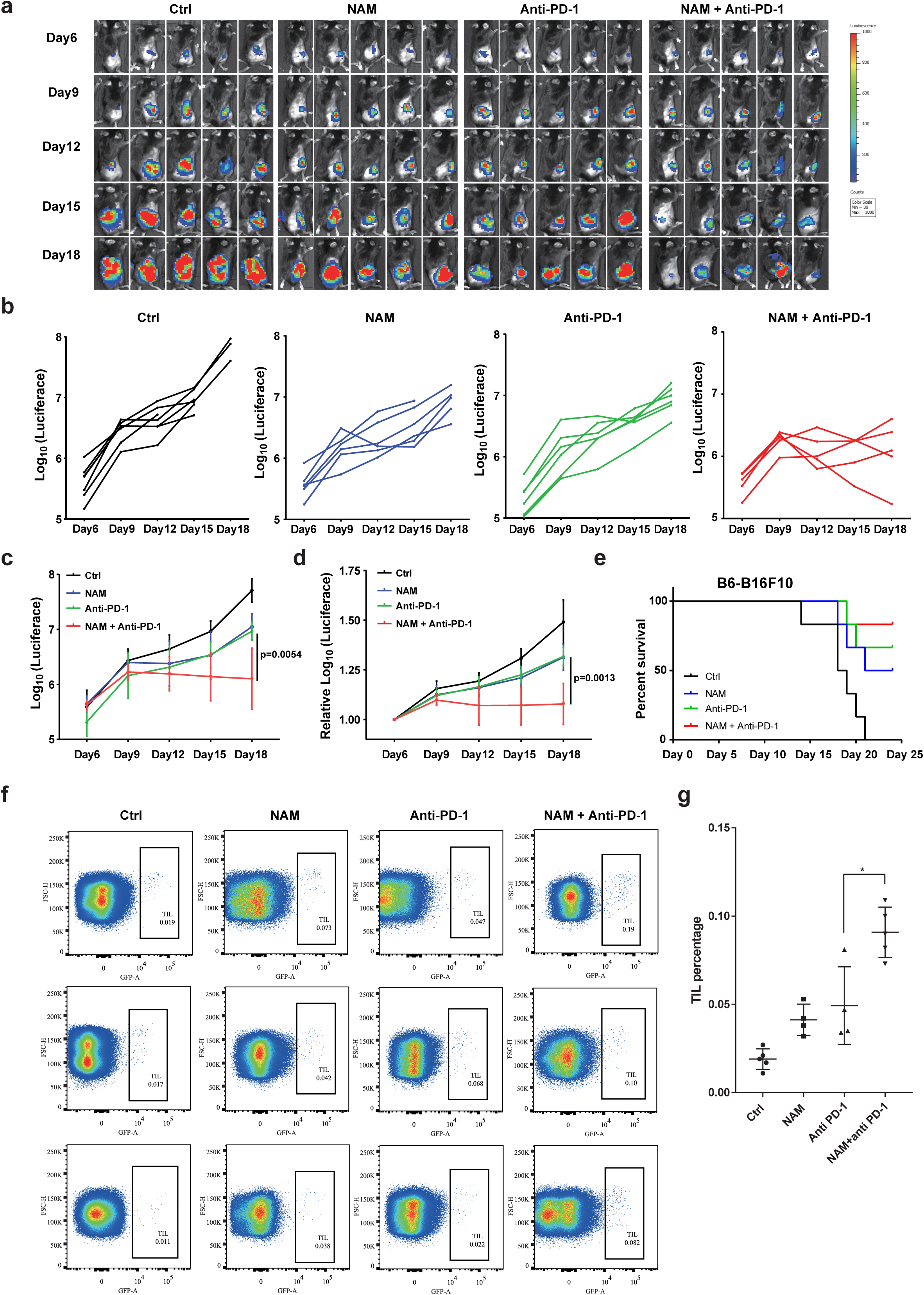
NAD^+^ supplementation enhances tumor killing function of anti-PD-1 treatment. **(a)** Bioluminescence imaging of B16F10 tumor model treated with indicated treatment (Ctrl, NAM, anti-PD-1, and anti-PD-1+NAM). n=5. **(b)** Tumor volume of individual mouse was calculated according to bioluminescence. n=5. **(c)** Tumor volume of each group was calculated according to bioluminescence. n=5. Data was shown as mean±SEM. **(d)** Relative tumor volume, normalized to initiate tumor volume, was calculated and plotted for each treatment group according to bioluminescence. n=5. Data was shown as ±SEM. **(e)** Log-rank test of survival curves. n=6. **(f)** Tumor infiltrated T cell ratio in each group was shown by FITC labelled anti-CD3 staining. **(g)** Statistical analysis of tumor infiltrated T cell percentage of each group. Data was shown as mean ±SEM.

There are three known sources of NAD^+^ in mammalian cells: *de novo* synthesis, and production via the Preiss–Handler (PH) or the salvage pathway ^39,49^. Recall our inhibitor screen and our NAD^+^ metabolism-related sgRNA CRISPR screening analyses indicating that NAD^+^ synthesis in T cells and especially in CD8^+^ cytotoxic T cells is apparently preferentially based on the NAMPT-regulated salvage pathway. We also discovered from the ovarian cancer patient samples and mouse melanoma tumors that TILs exhibit decreased intracellular NAD^+^ levels while the malignant tumor cells are reported to exhibit much higher NAD^+^ levels compared to normal cells ^33,50^. Interestingly, our conditional medium culture experiment also suggested that T cells showed lower NAD^+^ level when cultured with conditional medium, and this could be rescued by addition of exogenous NAD^+^ (Extended Data Fig. 12). We also observed that within tumor microenvironment, tumor cells harbor significantly higher concentration of NAD+ compared to TILs (Data not shown). We speculated that the nutrient competition in the hostile tumor microenvironment could be one of the extrinsic regulation factors of decreased NAD^+^ level in TILs, though this hypothesis still need further verification.

NAD^+^ is central to mitochondrial energy metabolism, and it also serves as a substrate for Sirtuin family members regulated deacetylation reaction and PARP family members mediated ADP-ribosylation ^36,37,51-53^. It was previously reported that a NAD^+^-Sirt1-Foxo1 axis controls the anti-tumor potential of hybrid Th1/17 cells generated from *ex vivo* cultures ^54^. In contrast to this role of NAD^+^ as a substrate for deacetylation and ADP-ribosylation, we here discovered that NAD^+^ can control T cell activation via regulation of energy metabolism. We found that CD8^+^ T cells, which are believed to be the main T cell population that functions in anti-tumor immunity, are especially sensitive to NAD^+^ limitation, a result emphasizing the major contribution of NAD^+^ to anti-tumor immunity. Consistently, we further established that NAD^+^ supplementation greatly elevates the anti-tumor function of T cells, an exciting result that immediately suggests the potential efficacy of such a supplementation strategy for improving immunotherapy outcomes generally.

## Acknowledgements

The authors thank the staff members of the Animal Facility at the National Facility for Protein Science in Shanghai (NFPS), Zhangjiang Lab, China for providing the support in mouse housing and care. The authors thank the staff members of the Multi-Omics Facility of Shanghaitech University for technical supports in LC-MS/MS experiment. The authors thank for Prof. Tiffany Horng’s critical reading and suggestions. The authors also thank for Li Chen, Shengmiao Chen, Jiali Zhang, and Wentao Li for their technical supports.

## Funding

This work was supported by the Ministry of Science and Technology of China (2018YFC1004603 to G.F.), the National Natural Science Foundation of China (31872831 to G.F.; 31670919 to H.W.), Science and Technology Commission of Shanghai Municipality (19JC1413800 to GF), the Shanghai Pujiang program (18PJ1407900 to GF), the Shanghai Shuguang Program (19SG55 to GF) and a ShanghaiTech University Startup grant (F-0202-17-041 to G.F.). H.W. is supported by 1,000-Youth Elite Program of China.

## Author information

YW, HW, and GF designed the study. YW, FW, YY, XX, QJ and SQ performed the experiments; LW, JC, and DL collected the human patient samples. XC and RW performed the isotropic labeling; YW, HW, and GF interpreted the data;YW and GF wrote the paper with the help from all authors; GF directed the whole project.

## Competing interests

Authors declare no competing interests.

## Supplementary Materials

Materials and Methods

Extended data figures 1-12

Extended data tables S1-S4

