## supplemental figure1-12 for "Potentiating the anti-tumor response of tumor infiltrated T cells by NAD^+^ supplementation"

^6^Lead Contact

^#^Equal Contribution

**This PDF file includes:**

Materials and Methods

Supplementary Text

Extended data figure1 to 12

Tables S1 to S4

**Materials and Methods**

**Cells and culture conditions**

293FT cells were cultured in DMEM (Gibco, 11965-092) supplemented with 10% FBS. Jurkat (JE6), JX003 (a single clone selected from Jurkat cells expressing Cas9) cells were cultured in RPMI supplemented with 5% FBS. Human T cells were cultured in RPMI (Gibco, 11875-093) supplemented with 10% FBS and 200U IL-2 (Novoprotein, GMP-CD66). Cells were maintained at 37°C in an atmosphere of 5% CO_2_. Lentivirus was produced using TrainsIT-2020 (Mirus Bio, MIR5405) based on the manufacturer’s instructions in the 293FT human embryonal kidney cell line.

The following compounds were purchased from the indicated suppliers and used at the indicated concentrations for in vitro studies in cell culture (unless otherwise stated): FK866 (Millipore, cat. no. 48-190-82), 10 nM to 1μM; NAM (Sigma, cat. no. N0636), 150μM; STK-118804 (Selleck, cat. no. S7316), 10 nM; NAD^+^ (Selleck, cat. no. S2518), 150μM; ATP (MCE, cat. no. HY-B2176), 5mM; Streptolysin O (Abcam, cat. no. ab126650), 0.5μg/ml; 2-DG (Selleck, cat. no. S4701), 5mM; CB-839 (Selleck, cat. no. S7655), 100nM.

Peripheral blood mononuclear cells were isolated by density gradient centrifugation and T cells were purified by Human T enrichment kit (Stemcell). Cells were activated with Dynabeads Human T-Activator CD3/CD28 (ThermoFisher) by 1:1 (beads : cell), cultured in RPMI-1640 (Gibco) supplemented with 5% FBS (Gemini Bioproducts) with 50Uml-1 human IL-2 (Recho) . 24 hours after activation, lentivirus was added into T cell cultures (MOI=5). 5 days after activation, the Dynabeads were moved. And, cells were expanded at the density of 0.5-1*10^6^ cells/ml in culture medium containing 200U/ml human IL-2. The CD19.41BBz CAR (FMC63) was constructed by linking sequences from a signal peptide derived from the human IL-2 gene to a CD19-specific scFv (FMC63), followed by a hinge and transmembrane domain from CD8, intracellular domains of 41BB and the intracellular domain of CD3-ζ were linked. Then, the sequence was subcloned into a pHR-EF1a based lentiviral backbone plasmid.

Medium used to culture K562-CD19 cells was collected and used as conditional medium. When the density of K562-CD19 cells reached 1 million/mL, passaged cells and collected the supernatant by centrifuging at 3,000 rpm for 15 minutes at 4 °C to remove cells and debris. Then the supernatant were filtered through 0.22 µm filter at 4 °C before using as conditional medium. For each experiment, the conditional medium was freshly collected.

K562-CD19-mCherry cells and anti-CD19-41BB CAR-T cells were used for co-culture experiment. 4 million K562-CD19-mCherry cells and 2 million anti-CD19-41BB CAR-T cells were mixed together and further cultured for 24 hours. Then cells were separated by flow cytometry and cell sorting (BD, FACSAriaⅡ).

**Whole genome sgRNA Library Screening**

For the genome-wide screens, the whole genome sgRNA plasmid library was established as previously reported . The library was ampliﬁed using Endura Electro Competent Cells following the manufacturer’s protocol (Endura, Cat #60242-1).

HEK293FT cells were seeded at 18 million cells in 15 cm poly-L-Lysine coated dishes 16 hours prior to transfection and cultured in DMEM + 10% FBS. Cells were transfected with the sgRNA transfer plasmids and 2nd generation lentiviral packaging plasmids, pMD2.G (Addgene, Cat #12259) and psPAX2 (Addgene, Cat #12260) using the Mirus transfection reagent (Mirus Bio, TransIT®-2020). The following day, media was refreshed. The viral supernatant was collected 48 hours post transfection and spun down at 300 g for 10 minutes, to remove cell debris. The virus was then concentrated by centrifugation at 27000 g for 120 minutes, at 4℃. Finally, lentiviral pellet was resuspended with RPMI and maintained at -80C.

Lentivirus was added directly to culture Jurkat cells at a 1:250 v/v ratio and gently mixed by tilting. Cells were expanded every two days, adding fresh media. On day 14, cells were stimulated with anti-TCR antibody (C305) and stained with anti-CD69. After gDNA isolation, sgRNAs were ampliﬁed and barcoded as in Joung et al. (2017), with adaptation to using a two-step PCR protocol. Post PCR, the samples were SPRI puriﬁed, quantiﬁed using the Qubit ssDNA high sensitivity assay kit (Thermo Fisher Scientiﬁc, cat #Q32854), and then analyzed on the 2100 Bioanalyzer Instrument. Samples were then sequenced on a HiSeq 4000 instrument (Illumina). Data were then analyzed and showed in Supplemental table S1.

The top2000 target genes of negative rank of enriched sgRNAs in the CD69^high^ group of whole genomic sgRNA screening in Jurkat and human T cells were used for overlapped candidates.

**Metabolic Inhibitors Screening**

Metabolic inhibitor library (Selleck, L5700) was purchased for this assay. Cells were pre-treated with inhibitors in 96-well plate for 24 hours. The Jurkat (JE6) cells were stimulated with anti-TCR antibody (C305) culture medium for 16 hours. The human T cells were stimulated with 3μg/ml anti-CD28 and 3μg/ml anti-CD3 for 16 hours. Cells were then stained with anti-CD69-APC, anti-CD25-PE and anti-ICOS-PE-Cy7. Data were shown in Supplemental table S2.

**T cell Activation Assay**

Jurkat cells were cultured at a concentration of less than 1 million/mL (for better stimulation) and were stimulated by a titrated anti-TCR antibody (C305). Sixteen hours after stimulation, cells were stained with anti-CD69-APC (Biolegend, cat. no. 310910) for 40min on ice. Then, the surface CD69 level was assessed using flow cytometry.

Human PBMC cells were cultured at a concentration of less than 1 million/mL. High-binding plate (Costar, cat. no. 3601) was coated with 3μg/ml anti-CD3 (eBioscience, cat. no. 13-0037-82) and 3μg/ml anti-CD28 ( [Biolegend, cat. no. 302914](http://www.baidu.com/link?url=au2AUBTjAF9eWMzyzdVDD8yUzcxrWRF6Qu6hLe85Q2hh-te7cJzZbMDItsz7BB44u_mzIRh1-yQ44w3WibIlCa)) at 4 ^o^C overnight. Then, the coated plate was washed with PBS before experiment. Human PBMC cells were stimulated by culturing in the coated plate. Sixteen hours after stimulation, cells were stained with anti-CD69-APC (Biolegend, cat. no. 310910), anti-CD25-PE (Biolegend, cat. no.302605) and anti-ICOS-PE-Cy7 (Biolegend, cat. no. 304303) for 40min on ice. Lastly, the surface levels of these markers were assessed using flow cytometry.

**NAD^+^ concentration measurement**

The NAD^+^ concentration was determined using a NAD^+^/NADH quantification colorimetric kit (Biovision, K337-100) according to the manufacturer’s standard protocol. NAD^+^/NADH assay kit is based on a alcohol dehydrogenase cycling reaction, in which the formed NADH reduces a formazan reagent. The intensity of the reduced product color, measured at 450 nm, is proportionate to the NAD^+^/NADH concentration in the samples.

Briefly, no less than 300,000 cells were washed with cold PBS and suspended in 200 μl chilly NADH/NAD extraction buffer. Total NADt was exacted by freeze/thaw two cycles (20min on dry ice, then 10min at room temperature). Vortex the extraction for 10 sec and centrifuge at 14000 rpm for 5 min. Transfer the extracted NADH/NAD^+^ supernatant into a new tube. To detect total NADt (NADH and NAD^+^), transfer 50 μl samples into 96-well plate for further reaction. To detect NADH, NAD^+^ needs to be decomposed before the reaction, by heating at 60℃ for 30min. Cool sample on ice. Transfer 50 μl of NAD^+^ decomposed samples into 96-well plate for further reaction.

Next, samples and NADH standards were incubated with NAD Cycling Mix at room temperature for 5 min to convert NAD^+^ to NADH. Add NADH developer for reaction 1 hours at room temperature. Read the plate at OD 450 nm. Then the NAD^+^ concentration could be calculated as NAD^+^=NADt-NADH. The readings were measured using SpectraMax i3 (MD).

**Ca^2+^ flux experiment**

Cells were diluted at a concentration of 1 × 10^6^ cells/ml in media. Add final concentration of 1.5 μM Indo-1 (Invitrogen, cat. no. I1226) into cells and incubate cells at 37°C for 30 minutes. After cells are labeled with Indo-1, wash 3 times with media and count. Resuspend cells at a concentration of 1 × 10^6^ cells/ml. For each sample, pre-incubate the cells with 2× C305 and 2 μM Ionomycin (Calbiochem, cat. no. 407953) at 37^o^C for 5 minutes prior to running on FACS. Then the Ca^2+^ was measured using flow cytometry.

**SgRNA Library Design and Cloning**

The NAD^+^ metabolism sgRNA library consists of 391 sgRNAs targeting enzymes participate in NAD^+^ synthesis and consumption, plus 3 NTCs, 3 sgZAP70 and 3 sgCBL as controls. All sgRNAs were designed against target sites that are of the format (N)_20_NGG using the website (https://portals.broadinstitute.org/gpp/public/analysis-tools/sgrna-design), and selected sgRNAs must pass the following off-targeting criteria: (i) the 11-bp seed must not have an exact match in any other promoter region, and (ii) if there is an exact off-target seed match, then the rest of the sgRNAs must have at least seven mismatches with the potential off-target site. After all sgRNAs that pass off-targeting criteria were generated, up to 5 sgRNAs per transcript were selected. All sgRNA sequences are shown in Supplemental table S3. The 20-nt target-specific sgRNA sequences were cloned into sgLenti sgRNA library vector MP783 (which is a gift from Prof. Wang Haopeng).

**Isotropic labelling**

Glutamine oxidation activity was determined by the rate of 14CO_2_ released from [U-14C]-glutamine . In brief, 1 million Jurkat cells were suspended in 0.5 ml fresh media. To facilitate the collection of 14CO_2_, cells were dispensed into 7ml glass vials (TS-13028, Thermo) with a PCR tube containing 50μl 0.2M KOH glued on the sidewall. After adding 0.5 μci [U-14C]-glutamine, the vials were capped using a screw cap with rubber septum (TS-12713, Thermo). The assay was stopped 2hr later by injection of 100μl 5N HCl and the vials were kept at room temperate overnight to trap the 14CO_2_. The 50μl KOH in the PCR tube was then transferred to scintillation vials containing 10ml scintillation solution for counting. A cell-free sample containing 0.5μci [U-14C]-glutamine was included as a background control.

Glycolytic activity was determined by measuring the detritiation of [5-3H]glucose . In brief, 1 million Jurkat cells were suspended in 0.5ml fresh media. The experiment was initiated by adding 1 µCi [5-3H] glucose and, 2hr later, media was transferred to a 1.5 ml microcentrifuge tube containing 50 μl 5N HCl. The microcentrifuge tubes were then placed in 20 mL scintillation vials containing 0.5 mL water with the vials capped and sealed. 3H_2_O was separated from unmetabolized [3H] glucose by evaporation diffusion for 24hr at room temperature. A cell-free sample containing 1 µCi 3H-glucose was included as a background control.

**Metabolite tracing and measurements**

For intracellular metabolite measurements, 1 × 10^7^ Jurkat cells were collected and washed twice using cold PBS. Metabolites were extracted using extraction solution (80% LC-MS-grade methanol, 20% Milli-Q water) with sonication on ice. Last, samples were centrifuged at maximum speed for 15 min at 4 °C and the supernatants were dried with vacuum and analyzed by LC–MS/MS on SCIEX [TripleTOF 6600 High Resolution Accurate Mass System](http://www.baidu.com/link?url=TVlCcTIsK4O9uzsU0V9v4e1kgJL1Uho4B3a60ySG_wG9hO3uaUdge98vOWFqU5MkJbES-gekm-PHnsGxAA-a9kPAJ4ZgRkBmo5mttiBUUf1h0Ui3S4V2hjoVx-caxROpepS_LVMZRr_FF_AA__Pxlq). Each group was replicated six times. The total metabolite results are shown in Supplemental table S4. Then MetaboAnalyst 4.0 was used for pathway enrichment analysis.

**ATP concentration measurement**

Cell Titer-Glo® (CTG) luminescent cell viability assay (Promega) was used to evaluate intracellular ATP level. Through reaction, the assay generates a luminescent signal proportional to the amount of ATP present, which also proportional to the number of viable cells.

In brief, cells were seeded and gradient diluted into a 96-well plate. Then, CTG reagent was added to each well and mixed for ∼15 min on an orbital shaker to induce cell lysis followed by luminescence reading. The signals were measured using SpectraMax i3 (MD). Results represent mean ± SEM from three independent experiments.

**ATP supplementation**

Cells were washed with calcium-free and magnesium-free 1 × DPBS to remove calcium and magnesium. Then cells were suspended with 450μl 1 × DPBS into a 1.5 ml centrifugal tube, treated with 50μl Streptolysin O (0.5μg/ml), and incubated 60 min at 37°C. Then cells were centrifuged at 400g for 4 min to discard the supernatant. Cells were re-suspend with 100μl 1 × DPBS, treated with 11μl 50mM ATP, and incubated in the 37°C for indicated time.

**Immunoblotting**

Cell extracts were prepared in 1% NP40 lysis buffer (20 mM Hepes pH 7.5, 150 mM NaCl, 1% NP40, 50 mM NaF, 1 mM Na_3_VO_4_, 10% glycerol, and protease inhibitor cocktail from Roche) at 4ºC for 30 minutes. Total protein concentration was determined using the BCA kit (Beyotime, P0010). An equal protein concentration was used for SDS–PAGE and transferred to a nitrocellulose membrane at 95 V for 90min at 4 °C. Following transfer, membranes were blocked using 5% nonfat milk in TBS/0.1% Tween 20 (TBST) for 1 h at room temperature. Membranes were incubated with primary antibodies overnight at 4 °C in 4% BSA/TBS + 0.025% sodium azide. Following incubation with the primary antibody, membranes were washed with TBST for 5 min at room temperature three times and then incubated with horseradish peroxidase-conjugated secondary antibodies (Cell Signaling Technology) made up in 5% nonfat milk for 1 h at room temperature. Following incubation with the secondary antibody, membranes were washed with TBST for 5 min at room temperature three times and then incubated with SuperSignal West Pico PLUS Chemiluminescent Substrate (Thermo Fisher) for visualization on film. The following antibodies were purchased from the indicated suppliers and used for immunoblotting (unless otherwise stated) at the indicated concentrations: anti-β-actin (Sigma, cat. no. A5441, clone AC-15); anti-NAMPT (Proteintech, cat. no.66385-1-lg); [Anti-Phosphotyrosine](http://supply.ncpss.org/wuzi/productOrder/viewProductDetail.action?productId=18386623) clone 4G10 (Millipore, cat. no.05-321); anti-pY493 Zap70 (Cell signaling technology, cat. no. 2704); anti-Zap70 (Cell signaling technology, cat. no. [2705](https://www.cst-c.com.cn/products/primary-antibodies/zap-70-99f2-rabbit-mab/2705?site-search-type=Products&N=4294956287&Ntt=zap&fromPage=plp)); anti-pZeta (Abcam, cat. no. ab68235); anti-Zeta(Abcam, cat. no. ab190728); anti-pErk (Cell signaling technology, cat. no. 9101); anti-Erk (Cell signaling technology, cat. no. 9102); anti-pY527 Src (Cell signaling technology, cat. no. 2105); anti-pY416 Src (Cell signaling technology, cat. no. 6943); anti-LCK (Cell signaling technology, cat. no. 2787;Biolegend, cat. no. LCK-01).

**Transmission electron microscopy**

Cells were fixed in 2.5% glutaralfehyde overnight at 4ºC and fixed in 2% osmic acid for 1.5 hours. Then cells were dehydrated with gradient ethano (5min in 30%, 5min in 50%, 15min in 70%, 15min in 80%, 15min in 95%, and 20min in 100%) and followed by 100% acetone dehydration for 20min. The cells were soaked in contain solution overnight at 4ºC. Epon 812 epoxy resin were used for cell embedding. Cell blocks were conserved in a 37 ºC drying oven. After ultrathin-section for no more than 70um, sections were stained with uranium acetate and lead citrate. The stained sections were used for electron microscopy analysis following a standard protocol at Institute of Biochemistry and Cell Biology, Chinese Academy of Science Imaging Core Facility. Sections were viewed on a JEM 1400 Transmission Electron Microscope (JEOL) operated at 120 kV and digital images were acquired with a Veleta 2K × 2K charge-coupled device camera (Olympus-SIS).

**Real-time oxygen consumption rate (OCR)**

The Seahorse XF24 Cell Culture Microplate (Agilent) was firstly coated with poly-lysine to increase cell affinity. Jurkat cells were plated at 8 × 10^5^ cells per well in a Seahorse XF24 Cell Culture Microplate (Agilent) with Agilent Seahorse XF Media (Agilent); a final volume of 525μl was placed in each well. To attach cells, cells were centrifuge at 200 × g (zero braking) for 1 minute. Cells were then incubated in a 0% CO_2_ chamber at 37 °C for 1 hr before being placed into a Seahorse XFe24 Analyzer (Agilent). For OCR experiments, cells were treated with 1μM oligomycin, 2μM carbonyl cyanide p-trifluoromethoxyphenylhydrazone (FCCP), and 0.5μM rotenone/antimycin.

**Intracellular Staining**

After co-cultured with target cells, cells were treated with intracellular protein transport inhibitor Brefeldin A (Selleck, cat. no. S7046) for 4 hours, to avoid potential protein secretion. Then cells were stained with [Zombie Violet](https://www.biolegend.com/en-us/products/zombie-violet-fixable-viability-kit-9341) (Biolegend, cat. no. 423113) for 20min on ice. For surface staining, the cells were incubated with antibodies for 20 min, washed, and fixed in 4% PFA for 15min at room temperature. After being washed with staining buffer (Biotech, FXP005), the cells were incubated with 0.15% Triton-100 for 10 min at room temperature. Then cells were stained intracellularly for 30 min, washed. All samples were read on [LSRFortessa](http://www.ndchina.com.cn/omscms.oms?id=643) cytometer (BD) and analyzed with FACS DIVA software v. 8.0 (BD Biosciences). Apc-anti-GzmB (Biolegend, cat. no. 372203) and Apc-anti-IFN-γ (Biolegend, cat. no. 502529) were used for staining.

***In vitro* killing assay**

Adjust the K562 cells at the concentration of 8×10^5^/ml; Mix K562-CD19-mCherry and K562-WT at 1:1 ratio (the final concentration of each K562 cell is 4×10^5^/ml). Then, adjust the concentration of CAR-T cells to 1.2×10^6^/ml. Then co-culture the CAR-T cells with mixed K562 cells at different ratio at 37 ^o^C for overnight. Harvest cells and stain cells with DAPI (1:200) before analyzing with flow cytometry.

After killing assay, we first gated K562 cell through cell size difference. Then we gated live cell with DAPI staining. Finally, we calculated the ratio of mCherry positive to evaluate the killing efficiency.

**Animal work**

Mice were raised in the animal facility of national center for protein science. All study protocols involving mice were approved by the Institutional Animal Care and Use Committee of the ShanghaiTech University and conducted in accordance with governmental regulations of China for the care and use of animals. In the subcutaneous injection model, 1 × 10^6^ K562-CD19-luciferace cells were suspended in 100 μL of PBS and subcutaneously injected into NSG mice. 4 days post subcutaneous injection, anti-CD19-41BB CAR-T cells were injected into mice via tail vein. Subcutaneous tumor growth was monitored periodically by injecting 150 μL of 10mg/kg D-luciferin (PerkinElmer) intraperitoneally and imaging the animal using a Xenogen imager (Xenogen IVIS-200 Optical *in vivo* imaging system). In the killing assay, mice were randomly divided into four groups: Ctrl, NAM, CAR-T and CAR-T+NAM. For mice in NAM and CAR-T+NAM groups, mice were treated with 500mg/kg NAM daily via intra-peritoneal injection. For mice in Ctrl and CAR-T groups, mice were treated with equal volume physiological saline daily via intra-peritoneal injection.

B6 mice were used for NAD^+^ concentration experiment. 6 weeks B6 mice were subcutaneously transplanted with 1x10^6^ B16F10 mouse melanoma cancer cells. Two weeks later, both tumors and spleen were harvest for sorting CD3^+^ tumor infiltrated (TIL) cells and CD3^+^ cells from spleen. Anti-CD3 (Biolegend, cat. no. 100235) was used for staining.

NSG mice were used for NAM supplementation experiment. 6 weeks NSG mice were injected with CAR-T cells via tail vein. Then mice were intraperitoneal injected with NAM or PBS daily. Three days later, spleen was harvest for sorting CD3^+^ T cells. Anti-CD3-APC (Biolegend, cat. no. 300412) was used for staining. Then the T cells were used for NAD^+^ concentration experiment.

B6 mice were used for anti-PD-1 and anti-CTLA-4 combination experiment. 6 weeks B6 mice were subcutaneously transplanted with 5x10^5^ B16F10 mouse melanoma cancer cells (anti-PD-1) or with 1x10^6^ MC38 mouse colorectal cancer cells (anti-CTLA-4). For anti-PD-1 treatment, mice were treated with anti-PD-1 ([BioXcell](http://www.nbs-bio.com/brand-40.html), cat. no. BE0146, 200 μg per mouse, dissolved in PBS) every three days (three times in total) six days after B16F10 inoculation. For anti-CTLA-4 treatment, mice were treated with anti-CTLA-4 (BioXcell, cat. no. BE0164, 10mg/kg, dissolved in PBS) twice every week (three weeks in total) six days after MC38 inoculation.

**Confocal microscopy**

Mouse tumor samples were harvest after palpable tumors formed, fixed in 4% PFA and dehydrated with 20% sucrose solution. After frozen dissection, Slides were washed with PBS and incubated with PBS with 3% BSA and 0.2% TritonX-100 for 30 min. Then, Slides were stained for biotin-labeled anti-CD3 antibody (eBioscience, cat. no. 13-0037-82) on ice overnight. The Slides were washed with PBS and stained with fluorescence secondary antibody (Invitrogen, 1:400) for 60 min at room temperature. Slides were mounted with Vectashield with DAPI and sealed with nail polish before image acquisition using a Zeiss LSM 880. Confocal images were analyzed and merged using ImageJ software.

**Patient samples**

7 ovarian cancer patient samples were collected with the approval by the institutional review committees of Nanjing Maternity and Child Health Care Hospital. 9 ovarian cancer patient samples were collected with the approval by the institutional review committees of the International Peace Maternity & Child Health Hospital of China welfare institute. Written informed consents were obtained from patients. The specimens were used for sorting CD3^+^ tumor infiltrated T (TIL) cells and CD3^+^ T cells from peripheral blood mononuclear cell (PBMC). Anti-CD3-APC (Biolegend, cat. no. 300412) was used for staining.

**Statistical Analysis**

Data were analyzed by Student's *t* test or Pearson correlation test if not mentioned in the text. *P* < 0.05 was considered as significant.

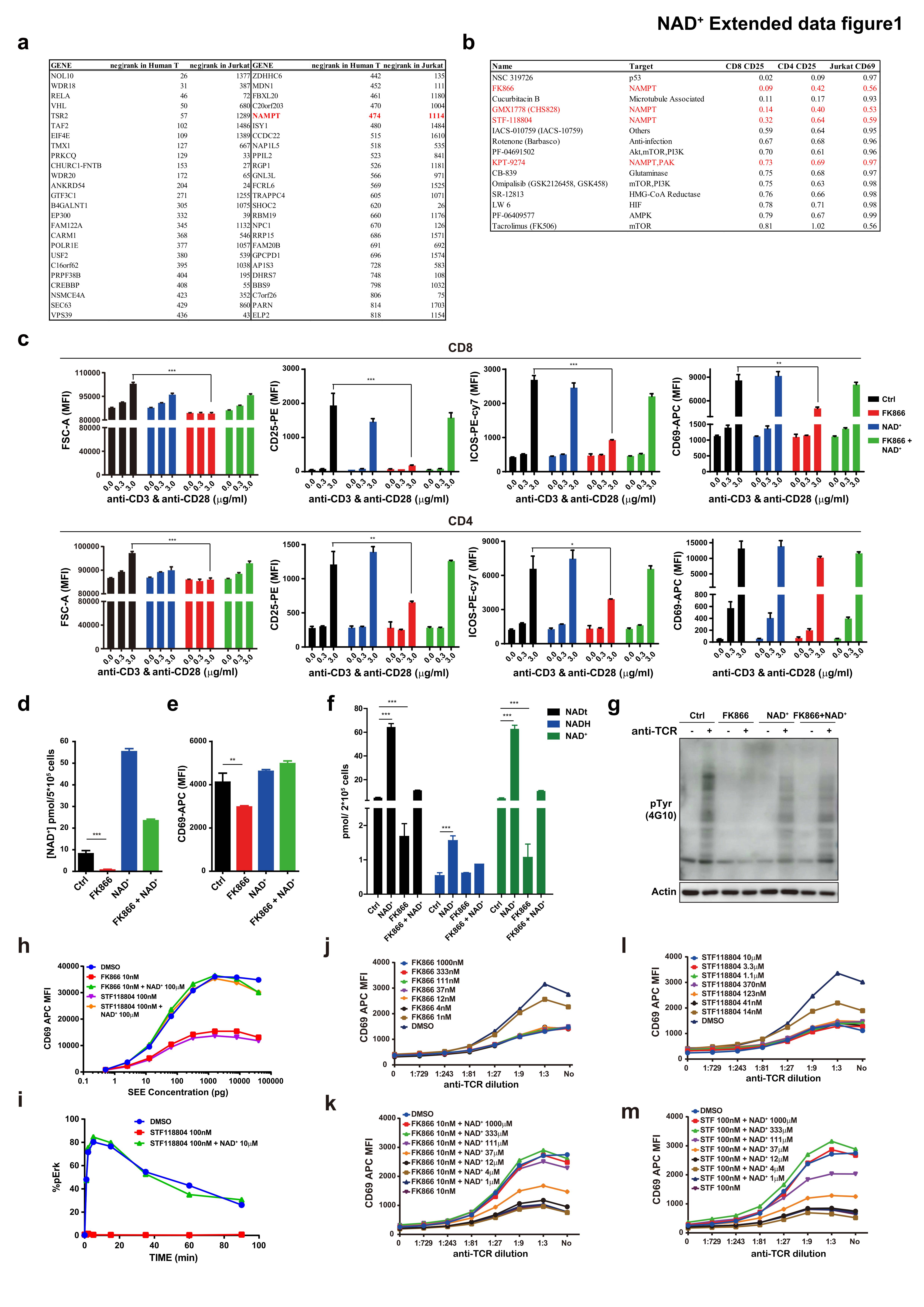

**Extended Data Figure 1 NAD^+^ is required for T cell function and generated mainly from salvage pathway in T cells. (a),** Whole genome sgRNA screens were performed in both Jurkat and primary human T cells to identify genes which are involved in T cell activation. List of top50 overlapped target genes from negative rank top2000 in both screens were illustrated. **(b)**, Metabolic inhibitor screenings were performed in Jurkat, human CD8^+^ and human CD4^+^ T cells. List of top inhibitors from the screen, as well as their matching target genes, was illustrated. **(c)**, Isolated human T cells in PBMC (both CD8^+^ and CD4^+^) were pre-treated as indicated (NAD**^+^**, FK866 or FK866 + NAD**^+^**) for 24 hours. Cells were then transferred to a plate coated with anti-CD3 and anti-CD28 antibodies for stimulation. Cell size and cell surface expression of CD25, ICOS, and CD69 were documented with statistic analysis. * p < 0.05, ** p < 0.01, *** p < 0.001. **(d)**, Human T cells from PBMC were first expanded by anti-CD3 and anti-CD28 stimulation, followed by NAMPT inhibitor FK866 treatment, as indicated. Concentration of NAD^+^ were measured and compared at different treating conditions. *** p < 0.001. **(e)**, Human T cells from PBMC were first expanded by anti-CD3 and anti-CD28 stimulation, followed by NAMPT inhibitor FK866 treatment, as indicated. Surface expression status of CD69 were measured and compared at different treating conditions. ** p < 0.01. **(f)**, Jurkat cells were pre-treated as indicated (NAD**^+^**, FK866 or FK866 + NAD**^+^**) for 24 hours. Concentration of NADt, NADH and NAD^+^ were measured and compared at different treating conditions. *** p < 0.001. **(g)**, Immunoblot of phosphotyrosine level in expanded T cells in human PBMC upon anti-CD3 and anti-CD28 stimulation. T cells were treated as indicated for 24 hours. After treatment, cells were rested in RPMI1640 without FBS for 1 hour. Then cells were stimulated with anti-CD3 (Biotin) for different time. **(h)** Cell surface expression of CD69 in Jurkat cells treated with NAD^+^ synthesis inhibitor (FK866 or STF118804) or NAD^+^ when activated with SEE. Jurkat cells were treated with indicated treatment for 24 hours and then cells were stimulated with SEE for 16 hours. **(i)** Phosphorylation level of ERK when stimulated with anti-TCR detected by FACS in Jurkat cells treated with STF118804. Jurkat cells were treated with indicated treatment for 24 hours. Then cells were rested in RPMI1640 without FBS for 1 hour. The cells were stimulated with anti-TCR, collected at indicated time point and were stained with phosphor-ERK. **(j to m)**, Surface expression of CD69 in Jurkat cells with indicated treatments. Jurkat cells were treated with indicated treatment for 24 hours. The cells were stimulated with anti-TCR.

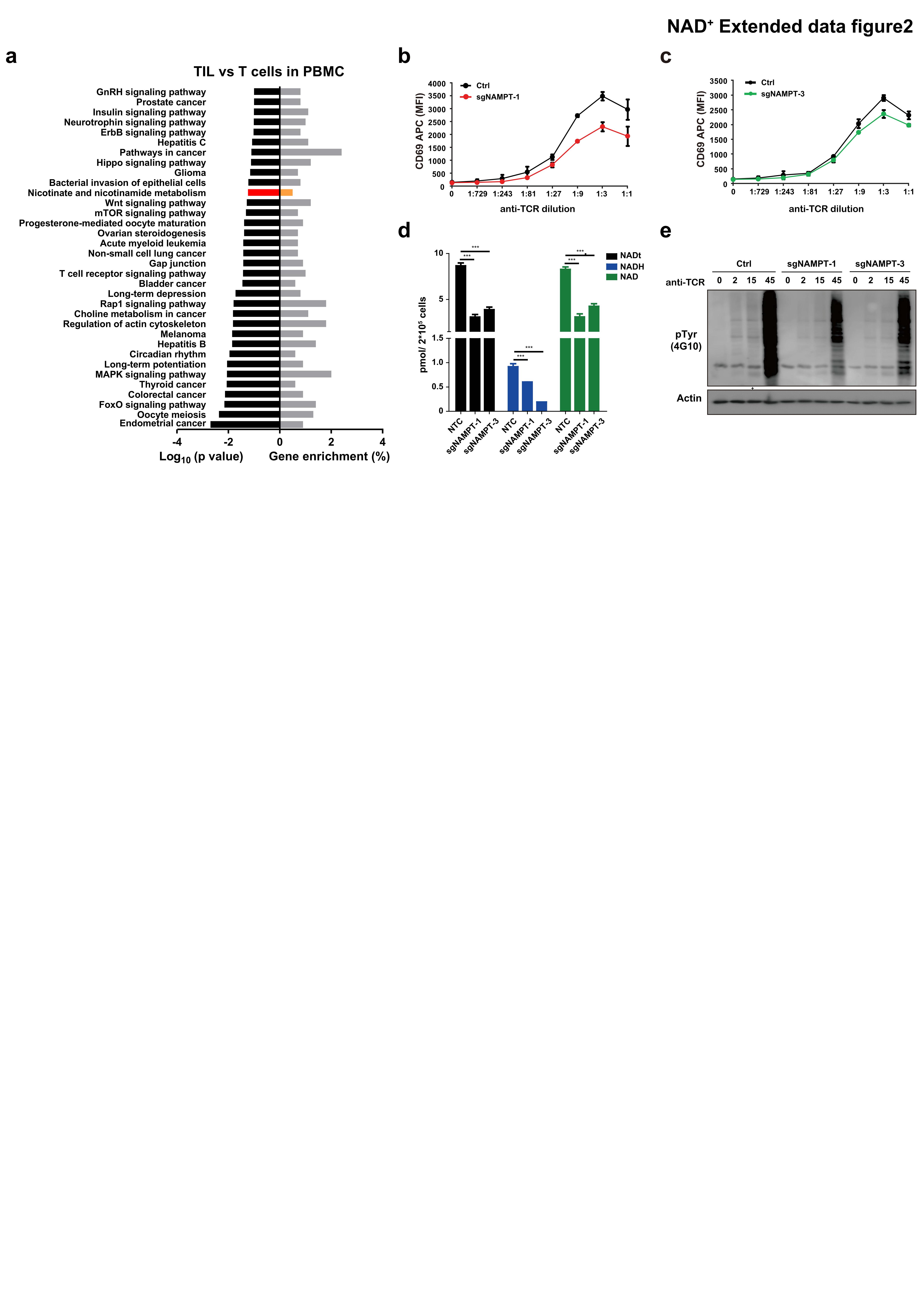

**Extended Data Figure 2 NAD^+^ deprivation restricts T cell activation. (a)**, Enrichment analysis with down regulated genes from public dataset (GSE42824). The genes with expression fold change (TIL/T) < 0.6 were used for enrichment analysis on DAVID website. **(b to c)**, JX003 cells were infected with virus expression indicated sgNAMPT, respectively. The titer of virus used for infection was calculated to infect about half of the Jurkat cells. After seven days, cells were stimulated with anti-TCR for 16 hours. Surface expression status of CD69 in Jurkat cells infected with sgNAMPT-1 (b) and sgNAMPT-3 (c) were illustrated. **(d)**, Concentration of NADt, NADH and NAD^+^ in Jurkat cells infected with sgNAMPT. JX003 cells were infected with virus expression indicated sgNAMPT, respectively. Cells were treated with Puromycin for selection. After seven days, cells were harvested for NADH concentration experiment. *** p < 0.001. **(e)**, Immunoblot of phosphotyrosine level in JX003 cells infected with sgNAMPT upon anti-TCR stimulation. Cells were treated with Puromycin for selection. Then, the cells were rested in RPMI1640 without FBS for 1 hour. After resting, the cells were stimulated with anti-TCR and collected at indicated time point for further immunoblotting analysis.

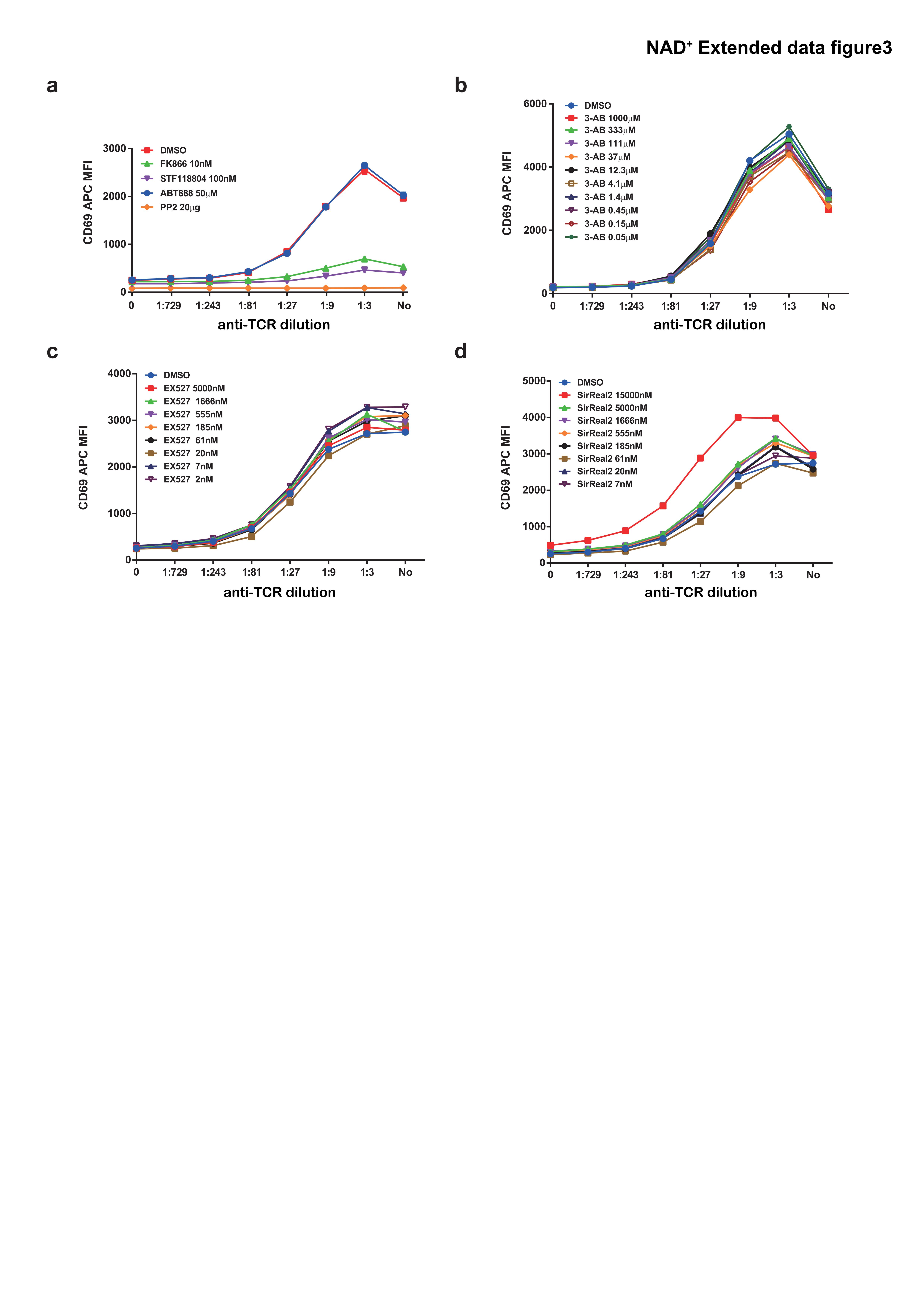

**Extended Data Figure 3 NAD^+^ regulate T cell activation via pathways independent of PARP mediated DNA repair or SIRT mediated histone deacetylation. (a)**, PARP1 specific inhibitor ABT888 had no effect on surface expression status of CD69 in Jurkat cells. Jurkat cells were pre-treated as indicated for 24 hours, followed by anti-TCR stimulation for 16 hours. Surface expression status of CD69 were measured and compared at different treating conditions. The inhibitor of SRC (PP2) was used as positive control. **(b)**, Pan-inhibitor of PARP family, 3-Aminobenzamide (3-AB) had no effect on surface expression status of CD69 in Jurkat cells. Jurkat cells were pre-treated as indicated for 24 hours, followed by anti-TCR stimulation for 16 hours. Surface expression status of CD69 were measured and compared. **(c)**, SIRT inhibitor EX527 had no effect on surface expression status of CD69 in Jurkat cells. Jurkat cells were pre-treated as indicated for 24 hours, followed by anti-TCR stimulation for 16 hours. Surface expression status of CD69 were measured and compared. **(d)**, SIRT inhibitor SirTeal2 had no effect on surface expression status of CD69 in Jurkat cells. Jurkat cells were pre-treated as indicated for 24 hours, followed by anti-TCR stimulation for 16 hours. Surface expression status of CD69 were measured and compared.

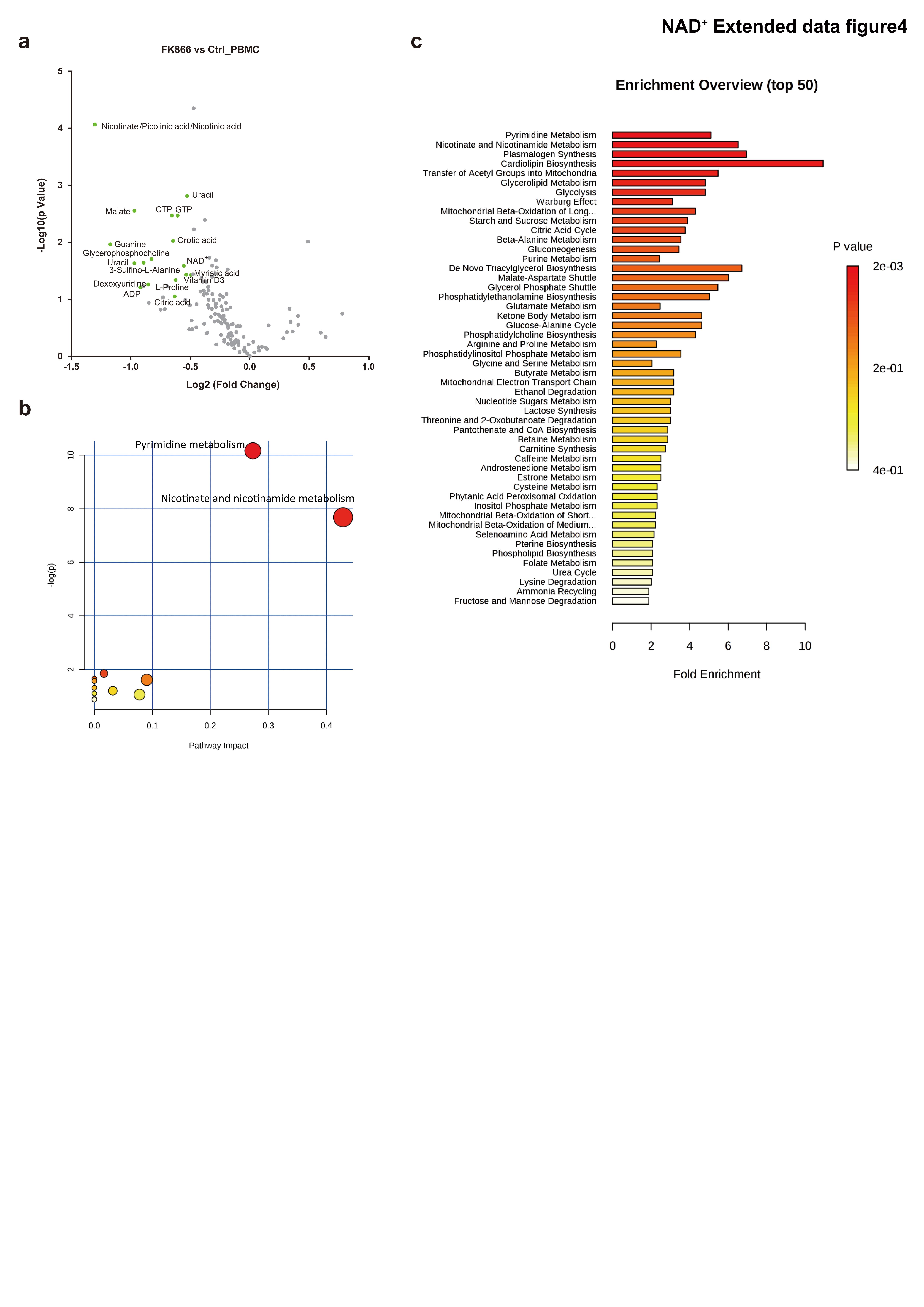

**Extended Data Figure 4 NAD^+^ regulates cellular energy metabolism through the TCA cycle in PBMC cells.** **(a)** LC-MS based metabolic profiling of PBMC cells treated with vehicle or FK866. 3*10^7^ PBMC cells, with or without 1μM FK866 co-treatment for 24 hours. Each group comprised three replicates. The green dots represent the metabolites down-regulated by FK866 treatment, and the red dots represent metabolites up-regulated by FK866 treatment. **(b-c)** Enrichment analysis of significantly changed metabolites. All significantly changed metabolites were used for metabolic pathway analysis with the MetaboAnalyst 4.0.

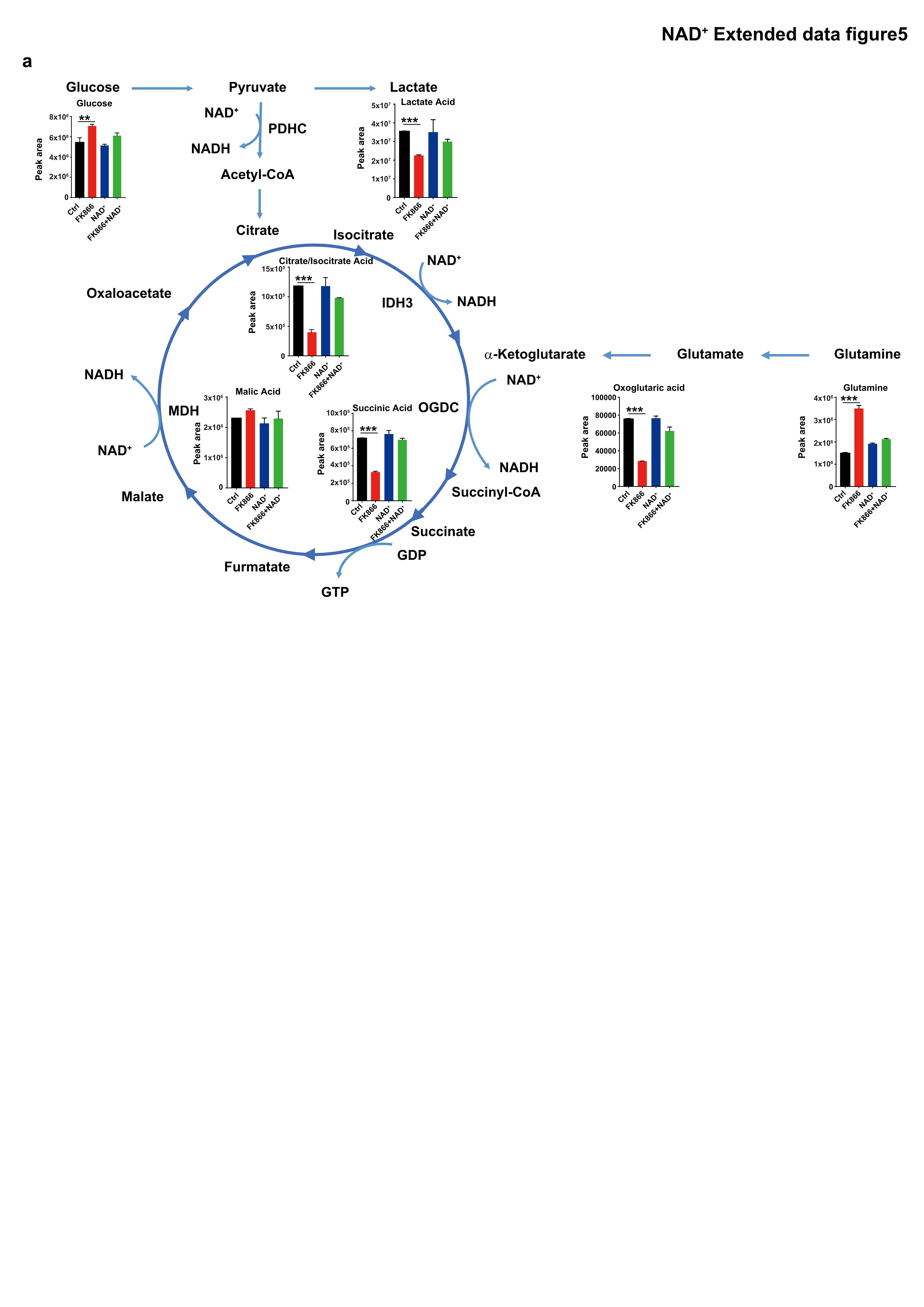

**Extended Data Figure 5 Metabolites in TCA cycle were decreased upon NAD^+^ depletion.** Metabolites related with TCA cycle detected by metabolic LC-MS in Jurkat cells. 1×10^7^ Jurkat cells were treated with indicated treatment for 24 hours. Then cells were harvested for LC-MS analysis. Each group was repeated six times. ** p < 0.01, *** p < 0.001.

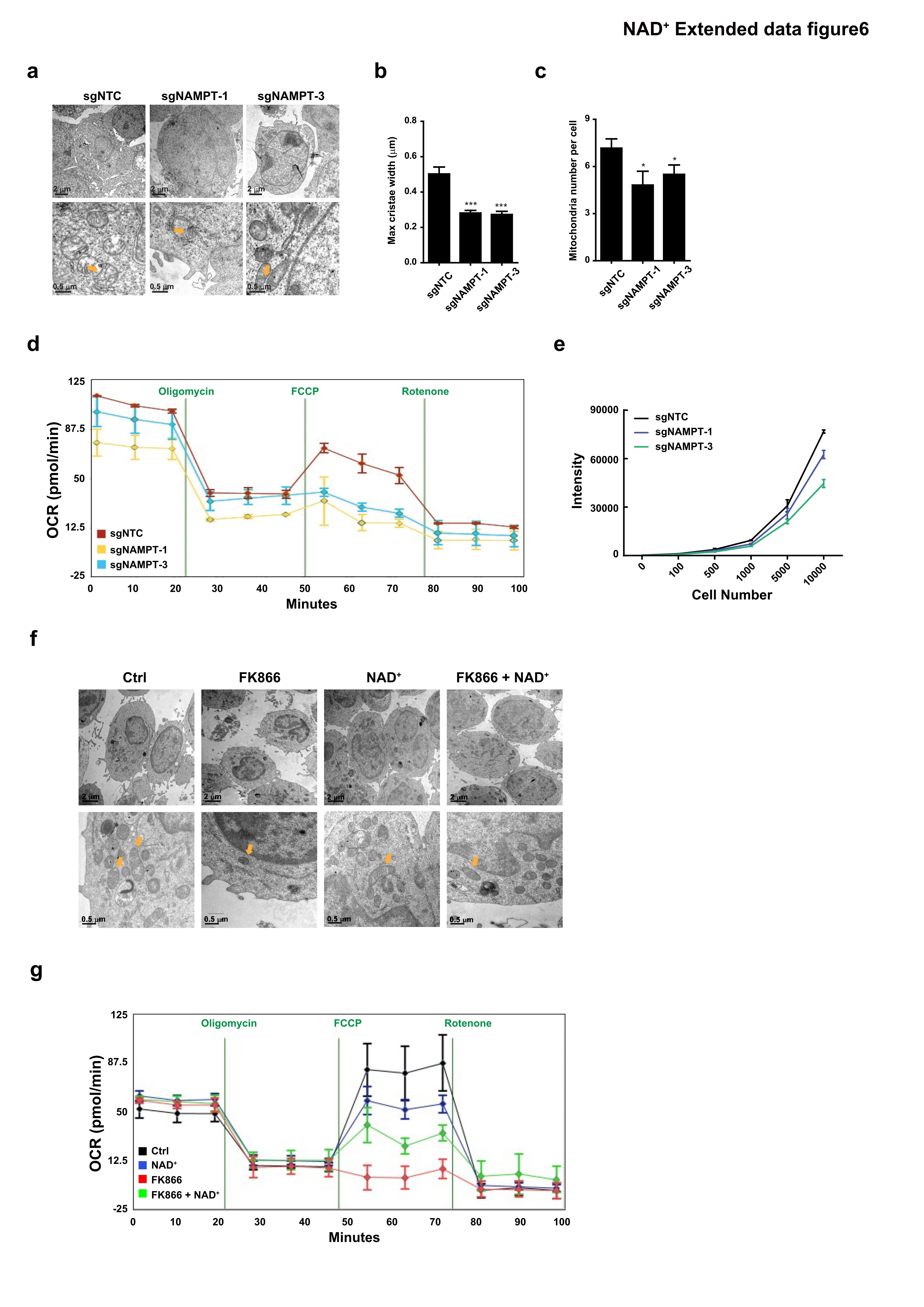

**Extended Data Figure 6 NAD^+^ regulates cellular energy metabolism. (a to c)**, Representable TEM images showed mitochondrial in JX003 cells (a) and statistic analysis (b-c). JX003 cells were infected with virus expressing indicated sgRNA (sgNTC or sgNAMPT), respectively. Cells were treated with Puromycin for selection. After selection, cells were harvested and prepared for TEM. * p < 0.05, *** p < 0.001. **(d)**, Oxygen consumption rate (OCR) under basal condition and in response to indicated mitochondrial inhibitors. JX003 cells were infected with virus expressing indicated sgRNA (sgNTC or sgNAMPT), respectively. Cells were treated with Puromycin for selection. After selection, cells were harvested and prepared for seahorse experiment. **(e)**, Cellular ATP level detected by CellTiter-Glo in JX003 cells infected with sgNAMPT. JX003 cells were infected with virus expressing indicated sgRNA (sgNTC or sgNAMPT), respectively. Cells were treated with Puromycin for selection. After selection, cells were harvested and prepared for CellTiter-Glo assay. **(f)**, Representable TEM images showed mitochondrial in PBMC cells. **(g)**, Oxygen consumption rate (OCR) under basal condition and in response to indicated mitochondrial inhibitors in PBMC cells.

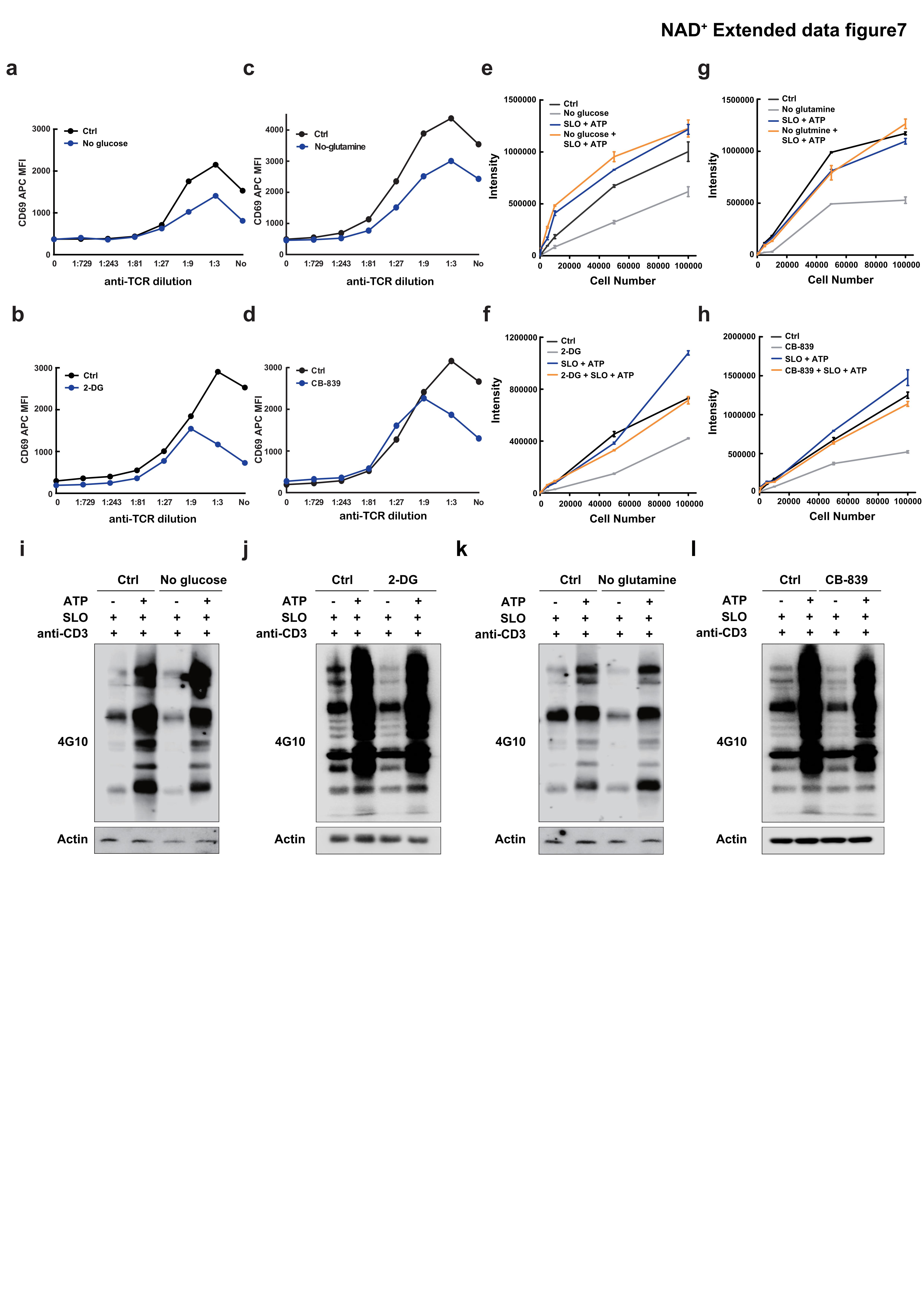

**Extended Data Figure 7 Mitochondrial metabolism regulates T cell activation. (a to b)**, Jurkat cells were treated with no glucose medium (a) or 2-DG (b) for 24 hours, followed by anti-TCR stimulation for 16 hours. Surface expression status of CD69 were measured and compared. **(c to d)**, Jurkat cells were treated with no glutamine medium (c) or CB-839 (d) for 24 hours, followed by anti-TCR stimulation for 16 hours. Surface expression status of CD69 were measured and compared. **(e to f)**, Cellular ATP level detected by CellTiter-Glo in expanded T cells cultured with no glucose medium (e) or 2-DG (f). Expanded T cells were treated with no glucose medium (e) or 2-DG (f) for 24 hours. Then the cells were harvested for CellTiter-Glo assay. **(g to h)**, Cellular ATP level detected by CellTiter-Glo in expanded T cells cultured with no glutamine medium (g) or CB-839 (h). Expanded T cells were treated with no glutamine medium (g) or CB-839 (h) for 24 hours. Then the cells were harvested for CellTiter-Glo assay. **(i to j)**, Immunoblot of phosphotyrosine level cultured with no glucose medium (i) or 2-DG (j). Expanded T cells were cultured with no glucose medium or 2-DG for 24 hours, followed by SLO treatment for 1 hour. Then cells were cultured in the absence or presence of ATP, with the stimulation of biotin labeled anti-CD3 for 2min and streptavidin (boosting the stimulation) for 5min. **(k to l)**, Immunoblot of phosphotyrosine level cultured with no glutamine medium k) or CB-839 (l). Expanded T cells were cultured with no glutamine medium or CB-839 for 24 hours, followed by SLO treatment for 1 hour. Then cells were cultured in the absence or presence of ATP, with the stimulation of biotin labeled anti-CD3 for 2min and streptavidin (boosting the stimulation) for 5min.

­­

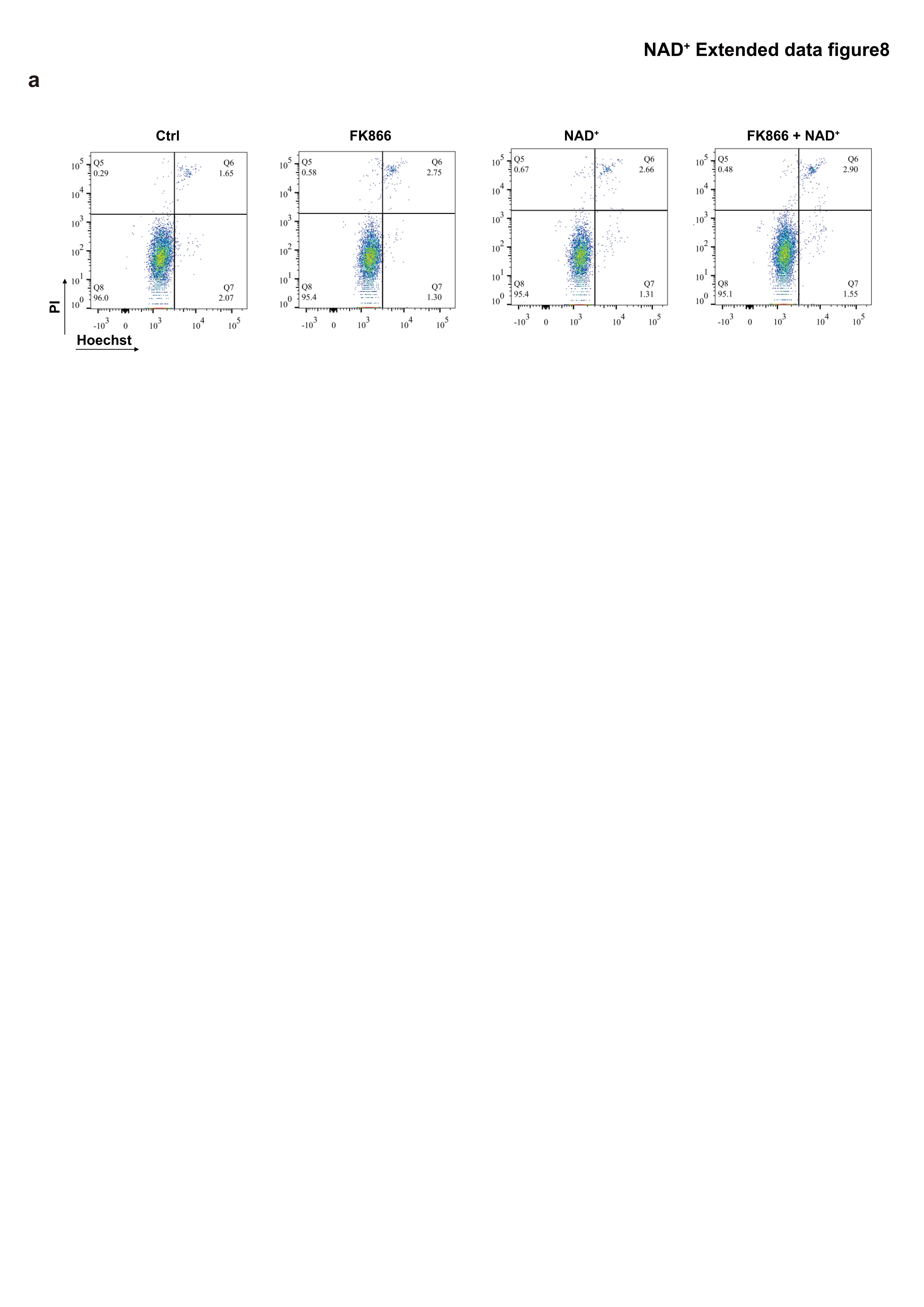

**Extended Data Figure 8 FK866 treatment did not induce cell death in CAR-T cells.** PI and Hoechst staining in 41BB CAR-T cells (a) with or without 1μM FK866 treatment for 24 hours.

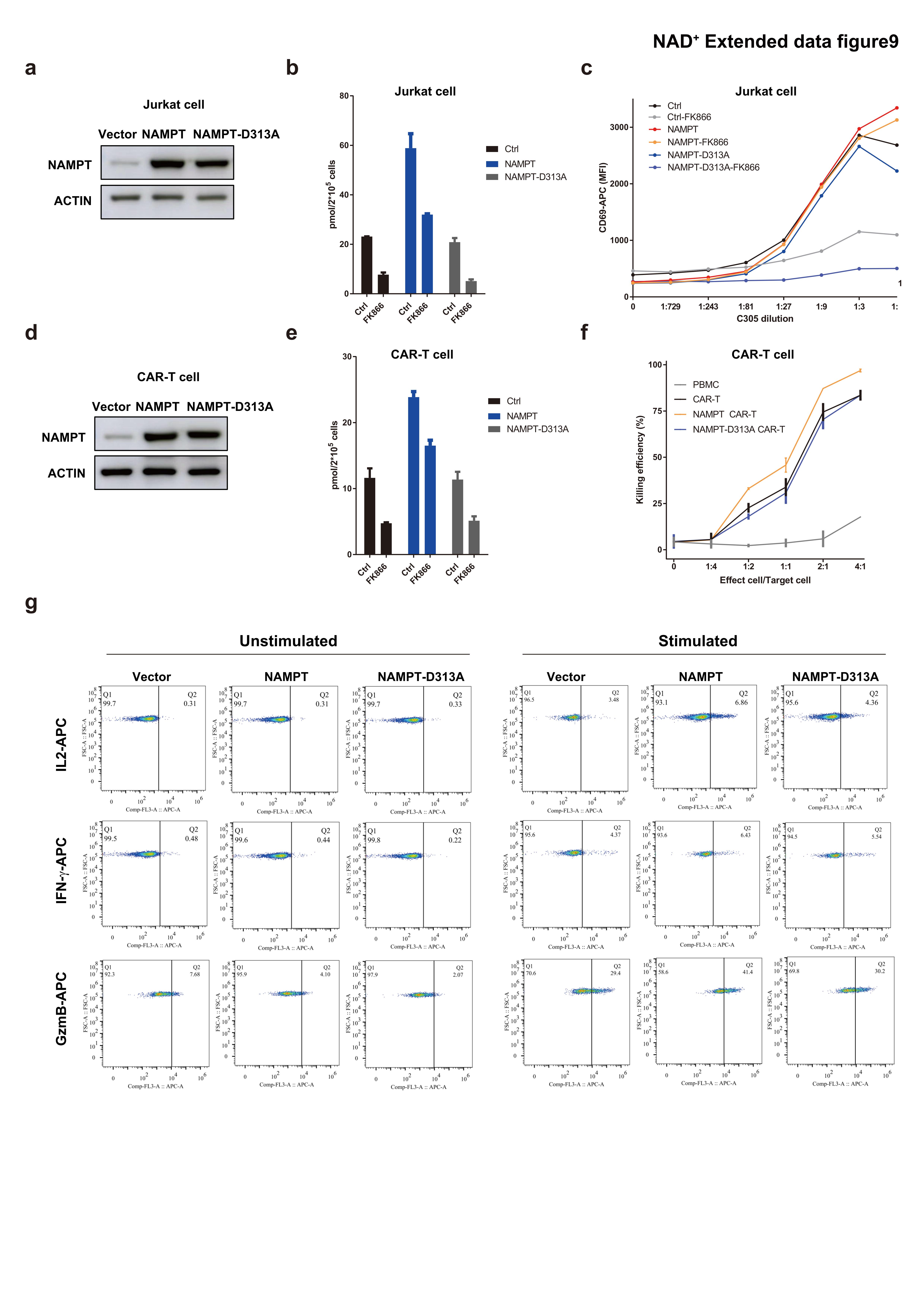

**Extended Data Figure 9 Overexpression of NAMPT could stimulate T cell activity. (a & d)**, WB showed NAMPT expression level in Jurkat T cells (a) and CAR-T cells (d) with or without NAMPT overexpression. **(b & e)**, NAD^+^ level in Jurkat T cells (b) and CAR-T cells (e) with or without NAMPT overexpression. **(c)** Jurkat T cells with or without NAMPT overexpression were pre-treated as indicated (FK866 or not) for 24 hours. Cells were then stimulated with anti-TCR (C305). Cell-surface expression levels of CD69 was measured using flow cytometry, respectively. **(f)** *In vitro* killing assay of target tumor K562-CD19 cells by anti-CD19-41BB CAR-T cells with or without NAMPT expression for 24hours. The killing efficiency was monitored by flow cytometry and calculated as the cell death ratio of the K562-CD19-mCherry cells. **(g)** Cytokine production, including GzmB, IFN-γ, and IL-2 of CAR-T cells with or without NAMPT expression.

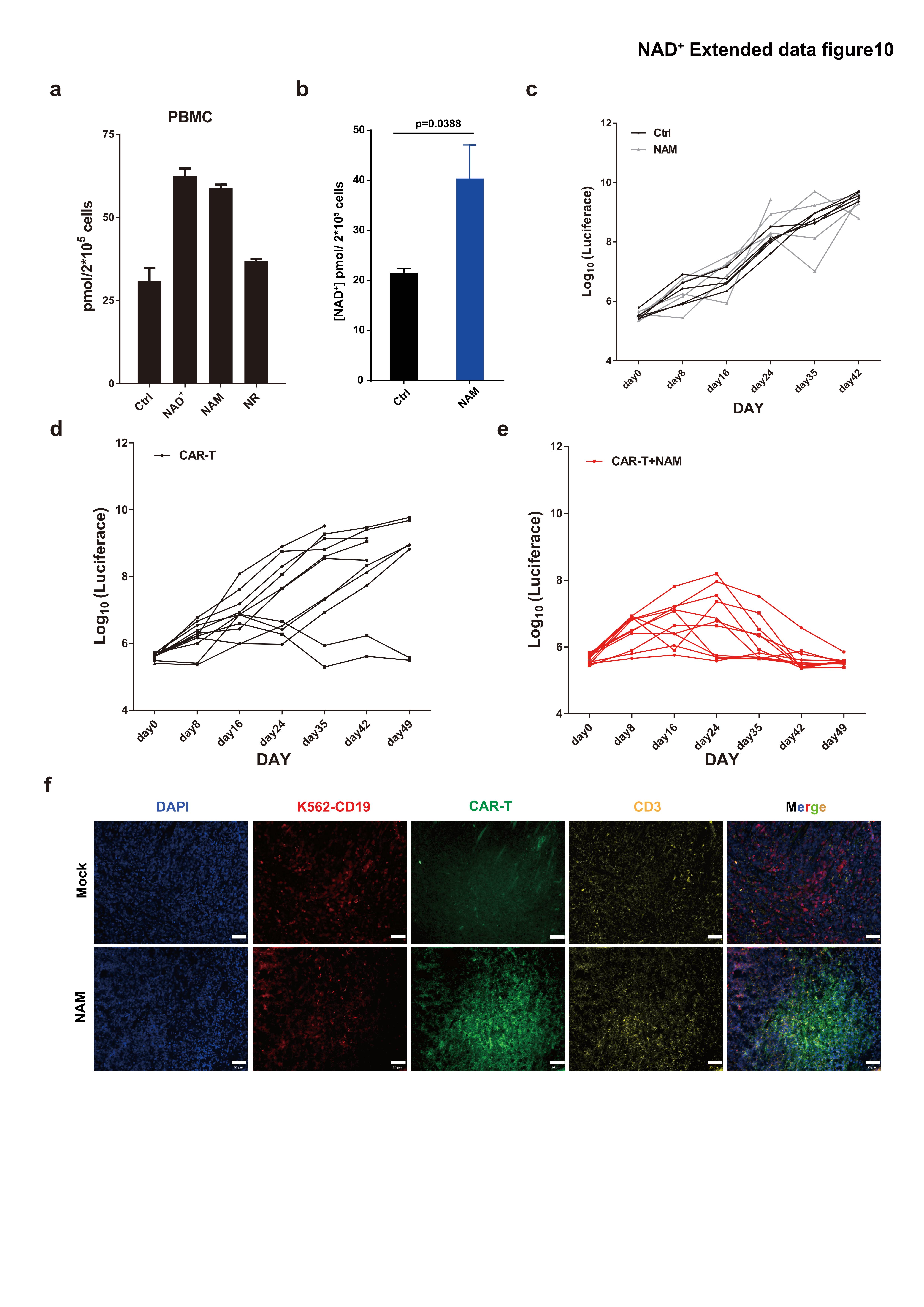

**Extended Data Figure 10 Restoring NAD^+^ concentration with NAM supplementation enhanced in vivo killing effect of CAR-T cell. (a)**, NAD^+^ concentration measurement of T cells in PBMC treated with different routes of NAD^+^ supplementation. Expanded T cells in PBMC were treated as indicated for 24 hours. Then, cells were harvested for NAD^+^ concentration experiment. **(b)**, Cellular NAD^+^ concentration detected in T cells in spleen from NSG mice transferred with CAR-T cells. The 6 weeks NSG mice were injected with CAR-T cells via tail vein and intraperitoneal injected with NAM for 3 days. Then the CD3^+^ T cells in spleen were isolated for NAD^+^ concentration experiment. **(c)**, Growth of tumor s.c. established K562-CD19-Luciferace upon adoptive transfer to NSG mice with or without NAM treatment. **(d)**, Ability of adoptively transferred anti-CD19-41BB CAR-T cells to control the growth of s.c.-established K562-CD19-Luciferace tumors in NSG mice. Tumor growth was measured by bioluminescence imaging. n=11. **(e)**, Ability of adoptively transferred anti-CD19-41BB CAR-T cells with NAM treatment to control the growth of s.c.-established K562-CD19-Luciferace tumors in NSG mice. Tumor growth was measured by bioluminescence imaging. n=12. **(f)**, Representable immunofluorescence images showed nuclei staining (DAPI), CD3 staining (YFP), CD19 (mcherry), and anti-CD19 CAR (GFP). Bar represents 50 μm.

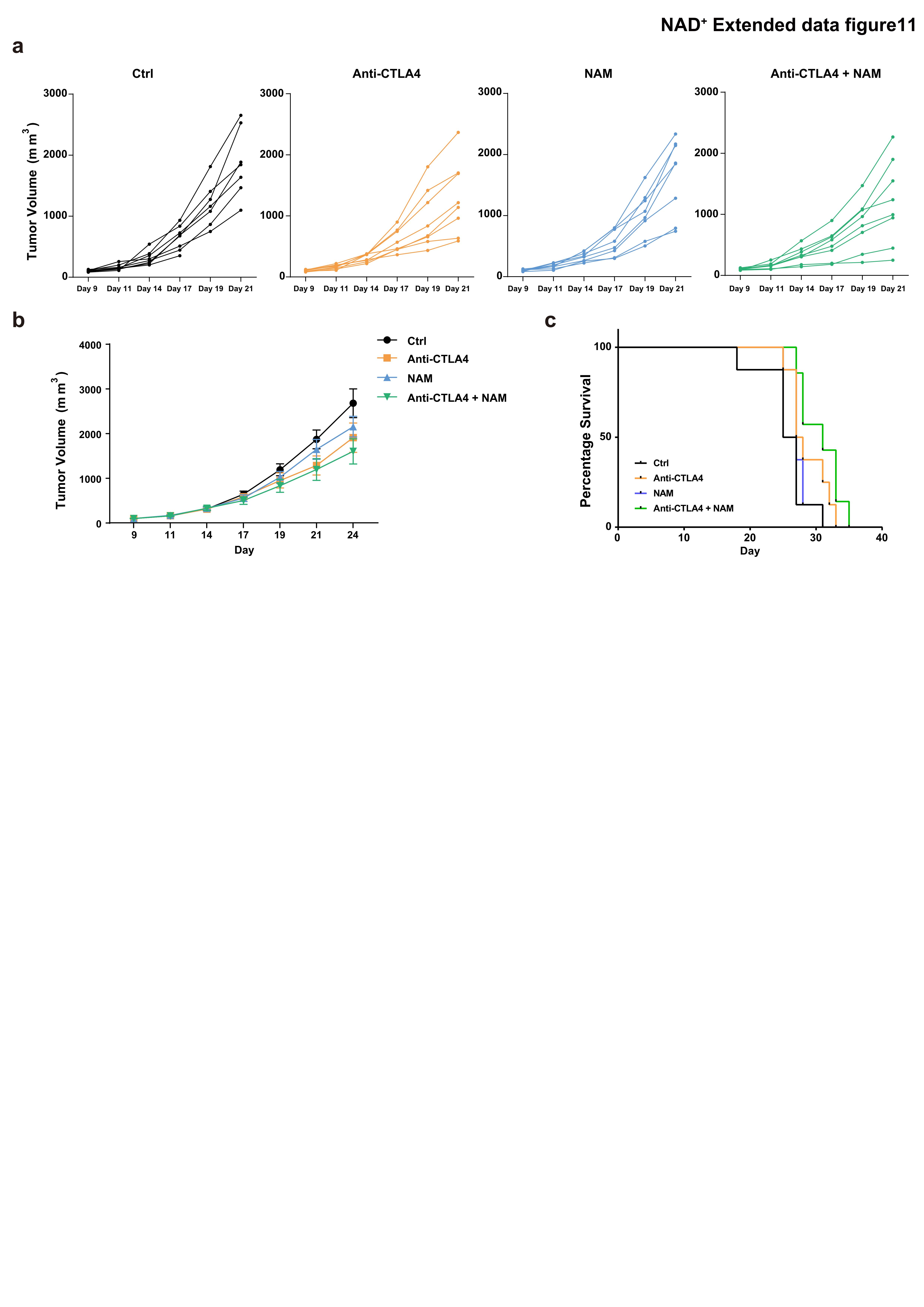

**Extended Data Figure 11 NAD^+^ supplementation enhances tumor killing function of anti-CTLA-4 treatment. (a)**, Tumor volume of individual mouse with indicated treatment (Ctrl, anti-CTLA-4, NAM, and anti-CTLA-4+NAM) was measured by Vernier caliper. **(b)**, Tumor volume of each group with indicated treatment (Ctrl, anti-CTLA-4, NAM, and anti-CTLA-4+NAM) was measured by Vernier caliper. Data was shown as mean±SEM. n=8. **(c)**, Log-rank test of survival curves. Ctrl group n=8, Anti-CTLA4 group n=8, NAM group n=8, and Anti-CTLA4+NAM group n=7.

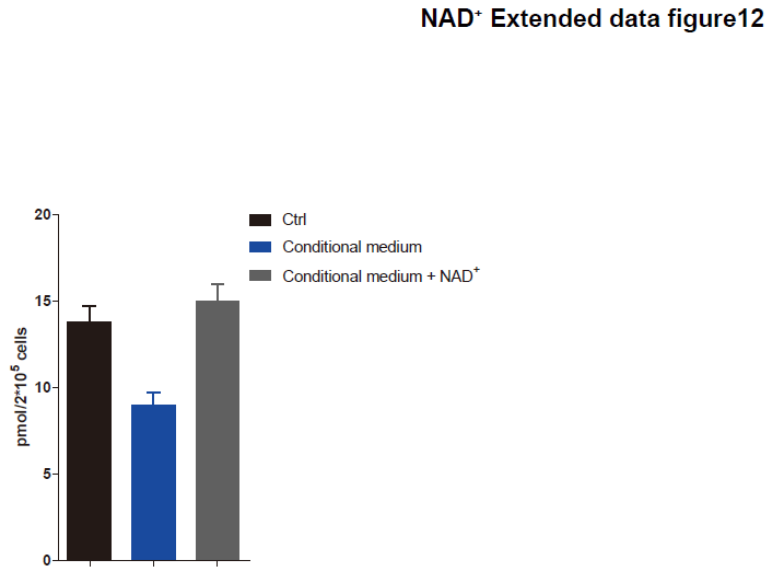

**Extended Data Figure 12** NAD^+^ concentration in CAR-T cells cultured with complete medium, conditional medium and conditional medium with NAD^+^ supplement. Data was shown as mean±SEM.
